# Supertertiary protein structure affects an allosteric network

**DOI:** 10.1101/2020.03.24.005553

**Authors:** Louise Laursen, Johanna Kliche, Stefano Gianni, Per Jemth

**Affiliations:** Department of Medical Biochemistry and Microbiology, Uppsala University, BMC Box 582, SE-75123 Uppsala, Sweden; Department of Chemistry-BMC, Uppsala University, Box 576, SE-751 23 Uppsala, Sweden; Istituto Pasteur-Fondazione Cenci Bolognetti and Istituto di Biologia e Patologia Molecolari del CNR, Dipartimento di Scienze Biochimiche “A. Rossi Fanelli,” Sapienza Università di Roma, 00185 Rome, Italy

## Abstract

The notion that protein function is allosterically regulated by structural or dynamic changes in proteins has been extensively investigated in several protein domains in isolation. In particular, PDZ domains have represented a paradigm for these studies, despite providing conflicting results. Furthermore, it is still unknown how the association between protein domains in supramodules, consitituting so-called supertertiary structure, affects allosteric networks. Here, we experimentally mapped the allosteric network in a PDZ:ligand complex, both in isolation and in the context of a supramodular structure, and show that allosteric networks in a PDZ domain are highly dependent on the supertertiary structure in which they are present. This striking sensitivity of allosteric networks to presence of adjacent protein domains is likely a common property of supertertiary structures in proteins. Our findings have general implications for prediction of allosteric networks from primary and tertiary structure and for quantitative descriptions of allostery.

## Introduction

Allosteric regulation is an essential function of many proteins, well established in multimeric proteins with well-defined conformational changes. From a biological perspective, such allostery plays important roles, for example in regulation of enzyme activity and binding of oxygen to hemoglobin, which was the basis for models such as MWC (1) and KNF (2). More recently the concept has been extended to monomeric proteins (3) and intrinsically disordered proteins (4, 5). Here, allostery is a process, where a signal propagates from one site to a physically distinct site, although the exact mechanism is elusive (6). Thus, as experimental approaches have developed the definition of the allosteric mechanism has evolved too and it is now spanning from the classical structure based allostery to the ensemble nature of allostery, and provides the framework for capturing allostery from rigid and structured proteins to dynamic and disordered proteins (7).

PDZ3 from PSD-95 has been extensively used as a model system for allostery in a single protein domain. Originally, it was used as a benchmark of a statistical method to predict allostery from co-variation of mutations (8) and has since then been subject to study by numerous methods (9). PDZ domains are small protein domains, around 100 amino acid residues, with a specific fold that contains 5-6 β strands and 2-3 α-helices (Figure 1C). The PDZ domain family is one of the most abundant protein-protein interaction domains in humans and it is often found in scaffolding or signalling proteins. PDZ domains bind to short interaction motifs (3-6 residues), usually at the C-terminus of ligand proteins. Looking at PDZ3 as a system to understand principles that underlie allosteric regulation reveals the complexity of the phenomenon. The first allosteric network in PDZ3 was determined by statistical coupling analysis and depicted a network that propagates from the binding pocket (α_2_ helix, β_2_ sheet and β_1_β_2_ loop) to the proposed allosteric site around the β_1_ and β_2_ strand. A part of the allosteric network was recently experimentally validated by a deep coupling scan (10) for the α_2_ helix in the PDZ3. However, their analysis was only applied to residues in direct contact with the peptide ligand. In fact, development of new approaches to capture allosteric networks in proteins have produced several distinct, but sometimes overlapping allosteric networks in PDZ3. Intriguingly, no two identical networks have been reported. The overall disagreement suggests that the resulting allosteric network is influenced by the choice of method to map it (9). A likely reason for the discrepancy is that allostery is caused by small energetic internal redistributions upon perturbation (11) in an overall cooperatively folded protein domain. It is clear that the context of a protein domain matters for its stability (12). Clearly, if the networks are so sensitive, it appears likely that the structural context of the protein domain would also affect any allosteric network.

**Figure 1:**
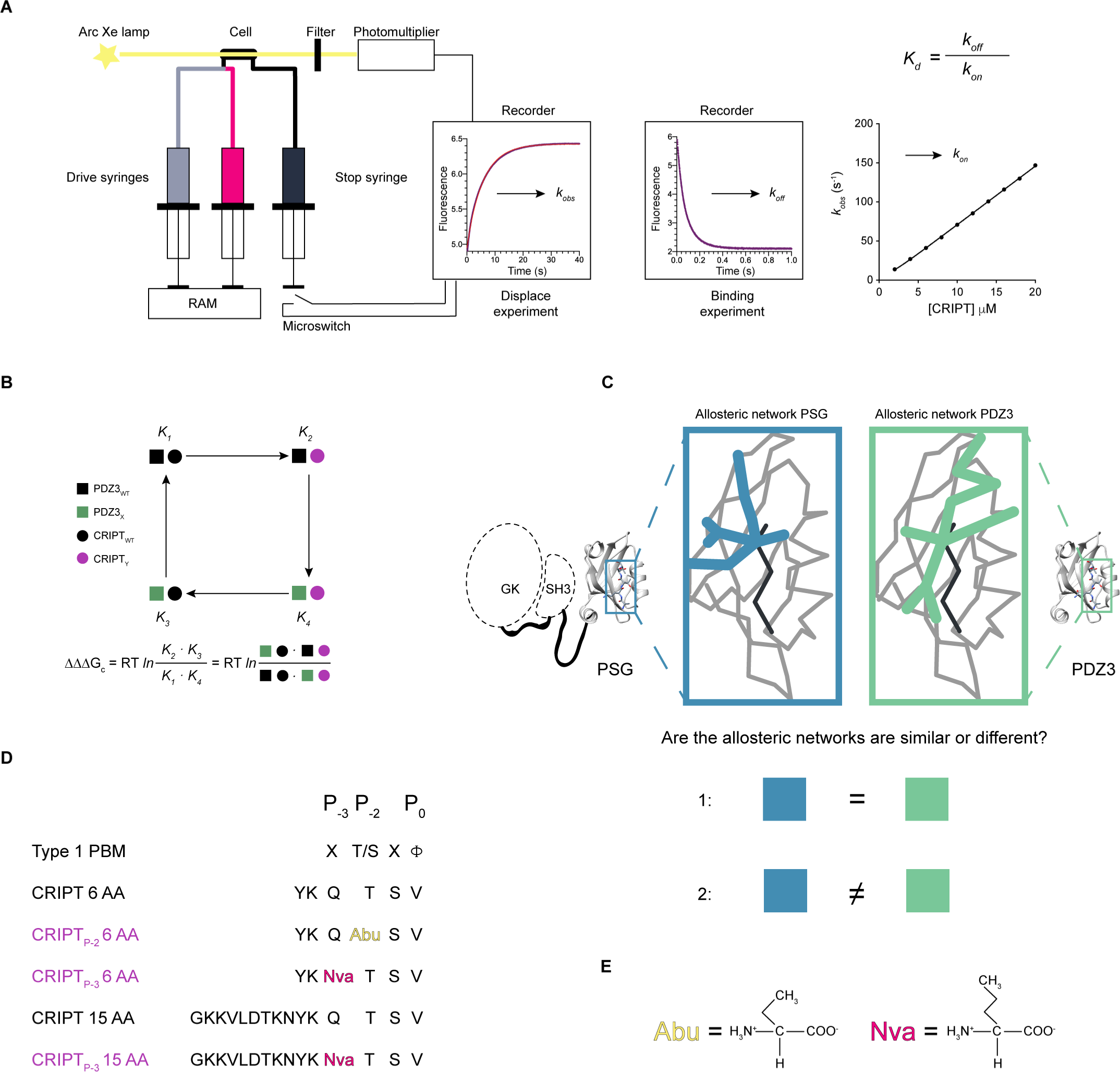
Illustration of experimental setup. A) Set-up for the stopped flow experiments and kinetic traces from displacement and binding experiments. *k*_obs_ values from binding experiments were plotted versus peptide concentration to obtain the association rate constant (*k*_on_), whereas the dissociation rate constant (*k*_off_) was obtained from displacement experiments. From these kinetic parameters, *K*_d_ values were calculated for each protein:peptide complex. B) Illustration of a double mutant cycle used to obtain the coupling free energy (ΔΔΔG_c_) between a residue in PDZ3 and in the CRIPT peptide ligand. C) Illustration of the overall question, to explore if the allosteric network in a single domain is dependent on the presence of supertertiary structure. To address the question, we compared the allosteric network within PDZ3 in the presence (blue) and absence (green) of the SH3 and GK domains. D) The different CRIPT peptides used in our experiments. Color codes: black represents WT CRIPT 6 AA or 15 AA and purple is CRIPT with a single mutation. A type 1 PDZ-binding motif (PBM) is defined by a hydrophobic residue at P_0_ and a Thr or Ser at P_-2_. The residues with mutated side chains are highlighted: P_-2_(Abu) and P_-3_(Nva). E) Sidechains for Abu (2-amino-butyric acid) and Nva (Norvaline).

Most PDZ domains are part of multidomain proteins. For example, PDZ3 is part of PSD-95, a scaffolding protein from the postsynaptic density that contains three PDZ domains, one SH3 and one guanylate kinase (GK) domain. PDZ1 and PDZ2 form a tandem domain supramodule whereas PDZ3, SH3 and GK form another supramodule, denoted PSG (Figure 1C), which is found in the whole MAGUK protein family to which PSD-95 belongs. Function and binding have been shown to be dependent on the interdomain architecture and dynamics, i.e., the supertertiary structure of the supramodules PDZ1-2 and PSG (13-15). Notably, while the allosteric networks for PDZ3 have been extensively studied, they have not been assessed in the context of the PSG supramodule. A few studies report a contribution to the allosteric network in PDZ3 from the α_3_ helix which connects PDZ3 with the SH3-GK tandem (16, 17) (Supplementary Figure 1). The potential role of the adjacent domains in the supramodule is particularly relevant since binding of protein ligands to the SH3-GK tandem is modulated by binding of cystein-rich interactor of PDZ3 (CRIPT) to PDZ3 (18).

To address the question of structural context for allosteric networks in single protein domains we mapped coupling free energies (ΔΔΔG_c_) upon peptide ligand binding to PDZ3 and PSG, respectively, using the double mutant cycle approach (Figure 1B) (19). A direct comparison of the mapped allosteric networks showed that similar patterns of energetic coupling were present around the binding pocket in both isolated PDZ3 and the PSG. However, the presence of the SH3-GK tandem resulted in strong coupling from the bound peptide ligand to the β_1_β_2_ loop, β_2_β_3_ loop and α_3_ helix that was not observed with the single PDZ3 domain. Our results show that the allosteric energy landscape of PDZ3 is highly sensitive to supertertiary structure in agreement with the conflicting previous studies on isolated PDZ domains.

## Results

We set out to test whether the allosteric network in PDZ3 is dependent on the supertertiary structure in which it is present, i.e., the interdomain organization and dynamics of the PSG supramodule (Figure 1C). To address this question and seek general principles that determine allosteric signal propagation in small protein domains we measured and compared coupling free energies (ΔΔΔG_c_) from mutational perturbations for PDZ3 and PSG, respectively. We also asked the following two questions for both PDZ3 and PSG: How is the allosteric network, as defined by the pattern of ΔΔΔG_c_ values, affected by the origin of perturbation (i.e., the residue, which is mutated in the peptide)? And how is the allosteric network affected by the length of the peptide ligand? We mapped the allosteric network using N-terminally dansyl labeled peptides derived from the ligand CRIPT (with either the last 6 amino acids or the last 15 amino acids, respectively, from CRIPT) with mutations at either position P_-2_ or P_-3_ (Figure 1D and E). The peptides used were CRIPT 6 AA (YKQTSV), CRIPT_P-3_ 6 AA (YKNvaTSV), CRIPT_P-2_ 6 AA (YKQAbuSV), CRIPT 15 AA (GKKVLDTKNYKQTSV) and CRIPT_P-3_ 15 AA (GKKVLDTKNYKNvaTSV). We could not obtain good data with CRIPT_P-2_ 15 AA (GKKVLDTKNYKQAbuSV).

### Experimental setup: Double mutant cycles as a method to map allosteric networks

In this study we applied double mutant cycles to quantify the intermolecular communication between residues in PDZ3 and CRIPT expressed as a coupling free energy, ΔΔΔG_c_ (Figure 1B). Double mutant cycle is a powerful approach (20, 21) and it has been applied to both intra- and intermolecular protein interactions (22-25). Our experimental setup was to measure kinetic rate constants for association (*k*_on_) and dissociation (*k*_off_) by stopped-flow experiments to deduce the affinity for the PDZ3:CRIPT interaction from the ratio *k*_off_/*k*_on_ (Figure 1A). Such rate constants have a high accuracy and precision, which results in relatively small errors in ΔΔΔG_c_. The largest experimental error is in the concentration of ligand, but the dansyl group used in the experiments provides accurate determination by absorbance.

The study was performed using our previously developed experimental system in which a Trp residue was engineered into PDZ3 (F337W) to enhance the change in fluorescence upon peptide ligand binding. F337W was previously shown to not affect the affinity of PDZ3 for the CRIPT interaction (26). In the present study we recorded circular dichroism spectra (200-260 nm) and determined the affinity by ITC for PDZ3 and PSG with and without the Trp337 probe (Supplementary Figure 2). The data showed that the F337W probe did not affect secondary structure or affinity and the pseudo wild types (containing F337W) are therefore referred to as WT PDZ3 and WT PSG.

In a double mutant cycle WT PDZ3, mutant PDZ3, WT CRIPT and mutant CRIPT were used to obtain four sets of *k*_*on*_ and *k*_*off*_ values for each coupling free energy. Thus, from the rate constants the affinities (*K*_*d*_ values) of the following four complexes were calculated: 1) PDZ3_WT_:CRIPT_WT_, 2) PDZ3_X_:CRIPT_WT_, 3) PDZ3_WT_:CRIPT_Y_, and 4) PDZ3_X_:CRIPT_Y_, where X and Y denote the respective mutation in PDZ3 and CRIPT (Figure 1B). The *K*_*d*_ values can be used to calculate ΔΔΔG_c_ between residue X in PDZ3 and residue Y in CRIPT. If the product of binding constants for single mutants equals the product of binding constants for WT and double mutant, then ΔΔΔG_c_ = 0 and no intermolecular interaction energy (or “communication”) exists between the probed residues X and Y. However, if ΔΔΔG_c_ ≠ 0 an intermolecular interaction is present between the two residues. We included a total of 32 mutations in PDZ3, 15 mutations in PSG, two mutations in CRIPT 6 AA and one mutation in CRIPT 15 AA, respectively. An example of a calculation is provided in Supplementary Figure 3.

In general we used conservative deletion mutations to conform to the assumptions underlying interpretation of mutations in proteins (27). Hence, residues in PDZ3 and PSG were substituted with Ala to prevent new interactions or steric clashes to be formed, the only exceptions being Ile→Val (also a conservative deletion mutation) and Gly→Ala. Circular dichroism spectra were recorded for all mutants to ensure intact secondary structure (Supplementary Figure 4). Binding kinetics (*k*_*on*_, *k*_*off*_) were studied for each mutant with all five peptide ligands included, i.e., CRIPT 6 AA, CRIPT_P-3_ 6 AA, CRIPT_P-2_ 6 AA, CRIPT 15 AA and CRIPT_P-3_ 15 AA (Figure 1D and Supplementary Figure 5). We chose unnatural amino acids to make the CRIPT mutations conservative, Thr→Abu and Gln→Nva, respectively. These mutations probe the effect of the functional groups on Thr (hydroxyl) and Gln (amide) without changing the length of the hydrocarbon chain (Figure 1E).

### Patterns of energetic coupling from two peptide positions overlap in the isolated PDZ3 domain

The allosteric network in PDZ3 was analyzed by mutating 32 out of 93 residues based on residue properties and previous studies. Thus, the mutations probed hydrophobic interactions (22) and charge properties (17), the role of the α_3_ helix (15), residues important for folding (28) and for dynamics of PDZ3 (29) (Figure 2, 3 and Supplementary Table 1). The coupling free energy for each position was mapped onto the structure of PDZ3 to visualize the allosteric network (Figure 2). Comparison of the coupling free energies reported by the two different peptide perturbations in CRIPT_P-2_ 6 AA (Thr→Abu) and CRIPT_P-3_ 6 AA (Gln→Nva), revealed that ΔΔΔG_c_ values are mainly positive and that the allosteric networks overlap (Figure 2, 3 and Supplementary Table 1). A positive coupling free energy means that the effect of the mutation is larger in WT PDZ3 than in the mutant, suggesting that WT PDZ3 is optimized for Thr in position P_-2_ and Gln in position P_-3_, when binding to CRIPT. Furthermore, we found a couple of residues unique for the respective allosteric network, e.g., position P_-3_ is coupled to F400 in the α_3_ helix and displays a different pattern of ΔΔΔG_c_ values in the β_2_β_3_ loop as compared to position P_-2_. However most of the reported energetically coupled residues are found within the proximity of the binding pocket, similarly to previous findings (22) (Supplementary Figure 6). In general, CRIPT_P-2_ 6 AA reports stronger residue coupling in comparison with CRIPT_P-3_ 6 AA in PDZ3. This is expected since the mutated hydroxyl group at P_-2_ forms a direct contact to the conserved His372 in PDZ3. Furthermore, the hydroxyl group in CRIPT_P-2_ 6 AA also has stronger energetic coupling to residues in PDZ3 compared with the γ-methyl group of residue P_-2_, mutated in a previous study (Thr→Ser, peptide denoted CRIPT_P-2(Ser)_ 6 AA) (22) (Supplementary Figure 6A and C). This result, while not surprising, clearly shows that the smaller and more specific the perturbation, the more precise is the mapping of coupling free energy and interpretation of allosteric propagation in PDZ3 and other proteins.

**Figure 2:**
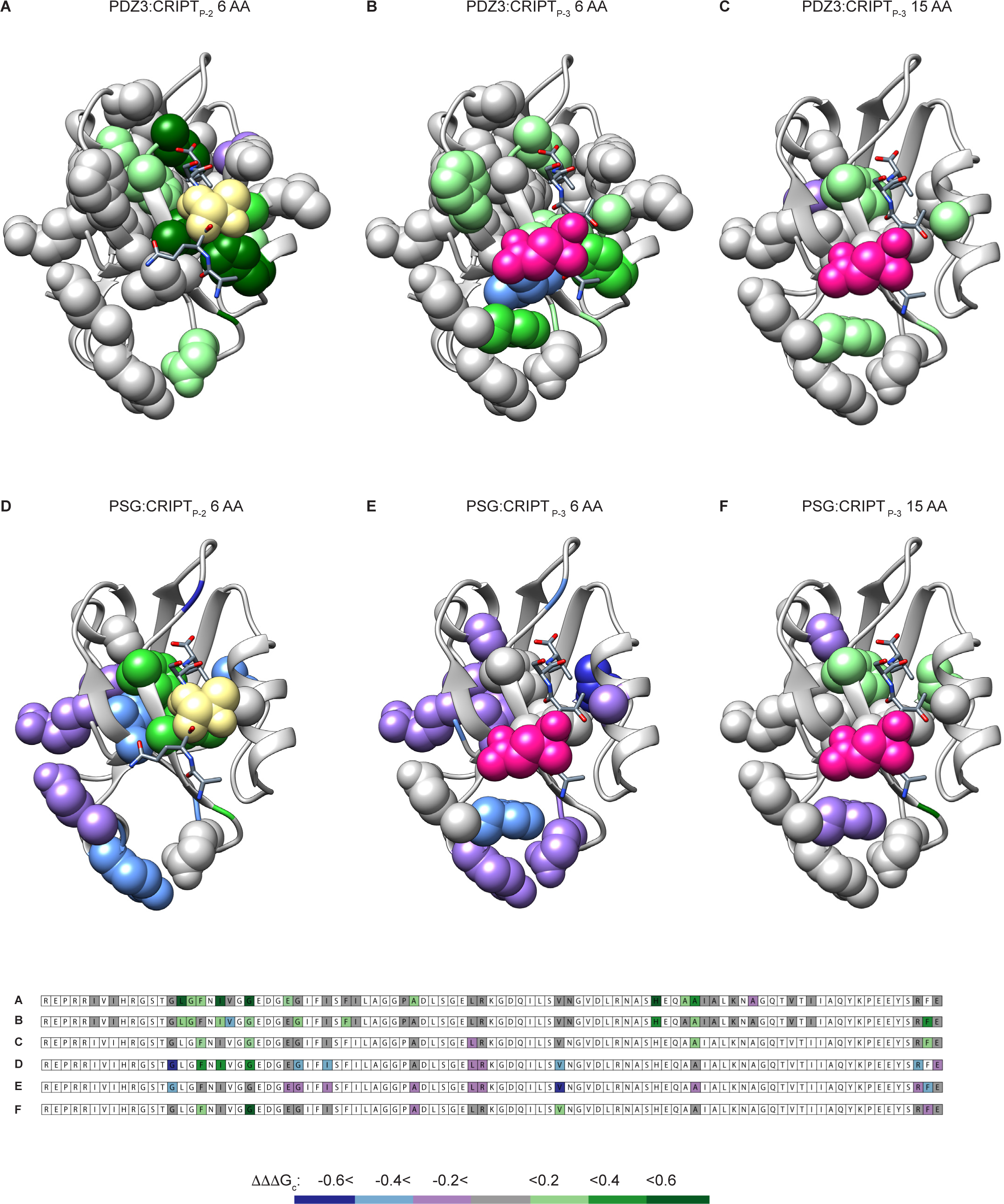
The allosteric network in PDZ3 is modulated by peptide sidechain, peptide length and presence of SH3-GK. The allosteric network, in terms of ΔΔΔG_c_ values mapped onto the PDZ3:CRIPT complex (pdb code: 1be9) for six different cases, where A-C display data on the isolated PDZ3 domain and D-F on the PSG supramodule. A and D) Propagation from CRIPT_P-2_ 6 AA (peptide perturbation Thr_-2_ → Abu) to residues in single domain PDZ3 (A) or PDZ3 in PSG (D). B and E) Propagation from CRIPT_P-3_ 6 AA (peptide perturbation Gln_-3_ → Nva) to residues in single domain PDZ3 (B) or PDZ3 in PSG (E). C and F) Propagation from CRIPT_P-3_ 15 AA (peptide perturbation Gln_-3_ → Nva) to residues in single domain PDZ3 (C) or PDZ3 in PSG (F). Color codes for mutations in CRIPT: yellow(P_-2_) and pink (P_-3_). Color code for the coupling free energy to side chains in PDZ3 (depicted with spheres): dark blue (< −0.6), light blue (< −0.4), purple (< −0.2), grey (−0.2 to 0.2), light green (>0.2), medium green (> 0.4) and dark green (>0.6). The primary structure of PDZ3 (residues 309-401) are shown below the structures and color coded based on coupling free energies.

**Figure 3:**
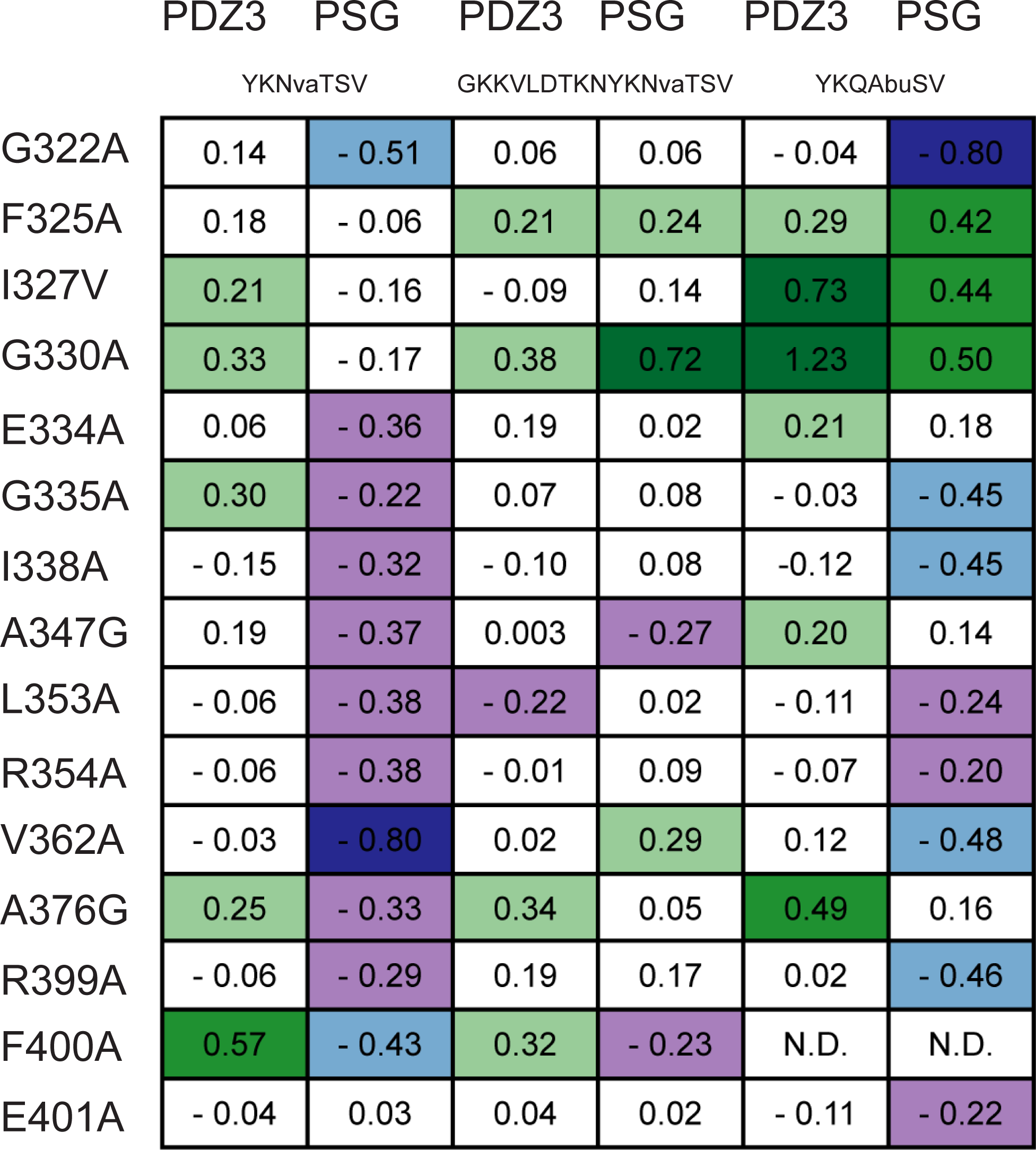
Coupling free energies (ΔΔΔG_c_) for sidechains in PDZ3 and PSG to sidechains in CRIPT. Values of coupling free energies mapped in Figure 2 are listed for 15 mutants in the six allosteric networks: PDZ3:CRIPT_P-3_ 6 AA, PSG:CRIPT_P-3_ 6 AA, PDZ3:CRIPT_P-3_ 15 AA, PSG:CRIPT_P-3_ 15 AA, PDZ3:CRIPT_P-2_ 6 AA and PSG:CRIPT_P-2_ 6 AA. Color code for ΔΔΔG_c_ values: dark blue (< −0.6), light blue (< −0.4), purple (< −0.2), grey (−0.2 to 0.2), light green (>0.2), medium green (> 0.4) and dark green (>0.6).

### The allosteric networks in PSG are distinct from those in isolated PDZ3

Binding of peptide ligands to PDZ3 can be analyzed by single exponential kinetics and is therefore well described by a simple two-state binding mechanism (22, 26). However, binding of CRIPT 6 AA to the PSG is double exponential suggesting a more complicated binding mechanism involving two distinct conformations of the PSG supramodule (30). We found previously that the two PSG conformations have similar affinity for CRIPT and showed that the fast kinetic phase, both in association and dissociation experiments, corresponds to one of the conformations (denoted PSG_A_). In the present study, this is the conformation we probe with regard to allosteric networks. In principle we could calculate ΔΔΔG_c_ values for PSG_B_ as well, but rate constants for the slow phase have larger errors (30). Thus, kinetic traces from experiments with CRIPT 6 AA and PSG were fitted to a double exponential function and *k*_*obs*_ values from the fast kinetic phases were used to estimate *k*_*on*_ and *k*_*off*_ from binding and displacement experiments, respectively (Supplementary Figure 5C and D and Supplementary Table 6-7).

15 mutations were chosen in PSG, covering different parts of PDZ3: the binding pocket (G322, F325, I327, E334), the common (G330, I338, A347, L353, V362, A376) and rare (G335, R354) proposed allosteric networks in PDZ3 (9) and the α_3_ helix (R399, F400, E401). Similarly to what we observed for PDZ3, the two allosteric networks in PSG, one determined for P_-2_ and the other for for P_-3_, overlap. However, comparison of PDZ3 and PSG reveals a completely different pattern of ΔΔΔG_c_ values for PDZ3 in the context of the PSG supramodule as compared with the isolated PDZ3 domain (Figure 2A-E and 3, Supplementary Table 1). Furthermore, while PDZ3 displays mainly positive coupling free energies PSG displays primarily negative values (Figure 2A-E and 3). Another striking difference is the strong coupling to α_3_ in PSG, which is not observed for PDZ3. Overall, PDZ3 has higher ΔΔΔG_c_ values around the binding pocket, whereas PSG has higher ΔΔΔG_c_ values among residues that are more globally distributed in the PDZ3 domain, e.g., in the β_1_β_2_ loop, β_2_β_3_ loop and α_3_ helix.

### The allosteric networks in PDZ3 and PSG are perturbed by longer CRIPT peptides

PDZ3 binds to a type 1 PDZ-binding motif (PBM) defined by a hydrophobic residue at position P_0_ (e.g., Val) and a hydrogen bond donor at P_-2_ (e.g., Thr) (Figure 1D). Residues outside the classical PBM influence the binding too, especially P_-5_ (Tyr) in the case of PDZ3:CRIPT (31, 32). However, beyond P_-5_, the affinity is not significantly affected by the length of CRIPT (33). However, it has not been reported if and how coupling free energies in PDZ and PSG are dependent on peptide ligand length. Therefore, we analyzed the effect of peptide length on the resulting allosteric network, as defined by ΔΔΔG_c_ values, for both PDZ3 and PSG. We report data for CRIPT_P-3_ 15 AA, but could not complete the analysis for CRIPT_P-2_ 15 AA since the kinetic phases were too fast for reliable stopped-flow experiments. Generally, the coupling free energies were weaker for CRIPT 15 AA compared with CRIPT 6 AA. The pattern was observed in both PDZ3 and PSG. Interestingly, the two networks contain both positive and negative coupling free energies, which could arise from lost selectivity and broader binding profile (34). Therefore the allosteric networks in PDZ3 and PSG upon binding of CRIPT 15 AA appear more similar (Figure 2C and F and 3). The weaker coupling free energy for CRIPT_P-3_ 15 AA is likely related to the fact that the dansyl in CRIPT 6 AA sits closer to the PSG. In fact, CRIPT 6 AA can sense the transition between PSG_A_ and PSG_B_, whereas CRIPT 15 AA cannot (30). Since we observe the pattern in both PDZ3 and PSG we can eliminate the concern that the five extra Trp residues in the SH3-GK tandem will disturb the F337W-dansyl signal pair. Thus, the data suggests that the weaker coupling free energy is a consequence of the 15 AA CRIPT. Thus, the probe influences the allosteric network. High coupling free energies are independent of perturbation origin, as CRIPT_P-2_ and CRIPT_P-3_ 6 AA report networks with similar coupling patterns, e.g., G322A and V362A for PSG and I327V, G330A and A376G for PDZ3. However, high coupling free energies seem to be affected by peptide length and adjacent domains. High negative coupling is reported for G322A and V362A in PSG:CRIPT_P-3_ 6 AA, but is not present for CRIPT_P-3_ 15 AA. The difference is also observed for PDZ3 and PSG, as G322A and V362A only show high negative coupling upon binding to CRIPT_P-2_ 6 AA in PSG. Thus, the probe and adjacent domains influence the allosteric network, underscoring the extraordinary sensitivity of the energetic connectivity among residues in this protein-ligand complex.

### Monotonic relationships between the allosteric networks

In an attempt to quantify any correlation among the coupling free energies in the different data sets we calculated Spearman correlation coefficients and plotted them in a heat map (Figure 4 and Supplementary Figure 7). We computed both Spearman and Pearson correlations, but choose to report the Spearman correlation as we cannot assume linearity and normal distribution between the variables. With the Spearman analysis and the associated heatmap we can visualize the allosteric networks (ranked coupling free energy values) that have a monotonic relationship. 14 (Figure 4) or 22 (Supplementary Figure 7) out of 32 mutations in PDZ3 were included in the correlation analysis to allow comparison with previously published data (22). Allosteric networks in PDZ3 were probed with four different CRIPT 6 AA peptides (Supplementary Figure 7A). None of the networks have a perfect correlation (r = 1), but all networks show a monotonic relationship (1 > r > 0), where several are significant (*) and not due to random sampling. The allosteric network probed with CRIPT_P-3_ 6 AA shows significant monotonic relationship to all networks, whereas CRIPT_P-2_ 6 AA only shows significant monotonic relationship to CRIPT_P-3_ 6 AA and CRIPT_P0_ 6 AA. Note that no relationship is seen between CRIPT_P-2_ 6 AA and CRIPT_P-2(Ser)_ 6 AA allosteric network, i.e., between that of the methyl group and that of the hydroxyl of Thr_-2_. This can be explained by the contribution from the hydroxyl group, which has a direct interaction to the sidechain of His372 (35) and underscores that the character of perturbation, not only residue, is important for the resulting allosteric network. Thus, a general Ala scanning would likely yield a distinct network for position −2 in the peptide. To further visualize correlations between allosteric networks, we quantified monotonic relationships by re-analyzing previous data from five sets of coupling free energy determined for SAP97 PDZ2 and an engineered circular permutated (cp) SAP97 PDZ2. Only one monotonic relationship was found for these data sets, between SAP97 PDZ2:GluR-A_P-2_ 9 AA and cp SAP97:PDZ2 GluR-A_P-2_ 9 AA (25, 36) (Supplementary Figure 7B). The authors highlighted that stronger coupling free energies were reported for PDZ3 as compared with the structurally similar SAP97 PDZ2 and cp SAP97 PDZ2, thus suggesting that side chains rather than backbone govern the magnitude of the allosteric network (25). This is further supported by our data for PDZ3 with CRIPT_P-2_ 6 AA and CRIPT_P-2(Ser)_ 6 AA (Supplementary Figure 6A and C and Supplementary Figure 7A).

**Figure 4:**
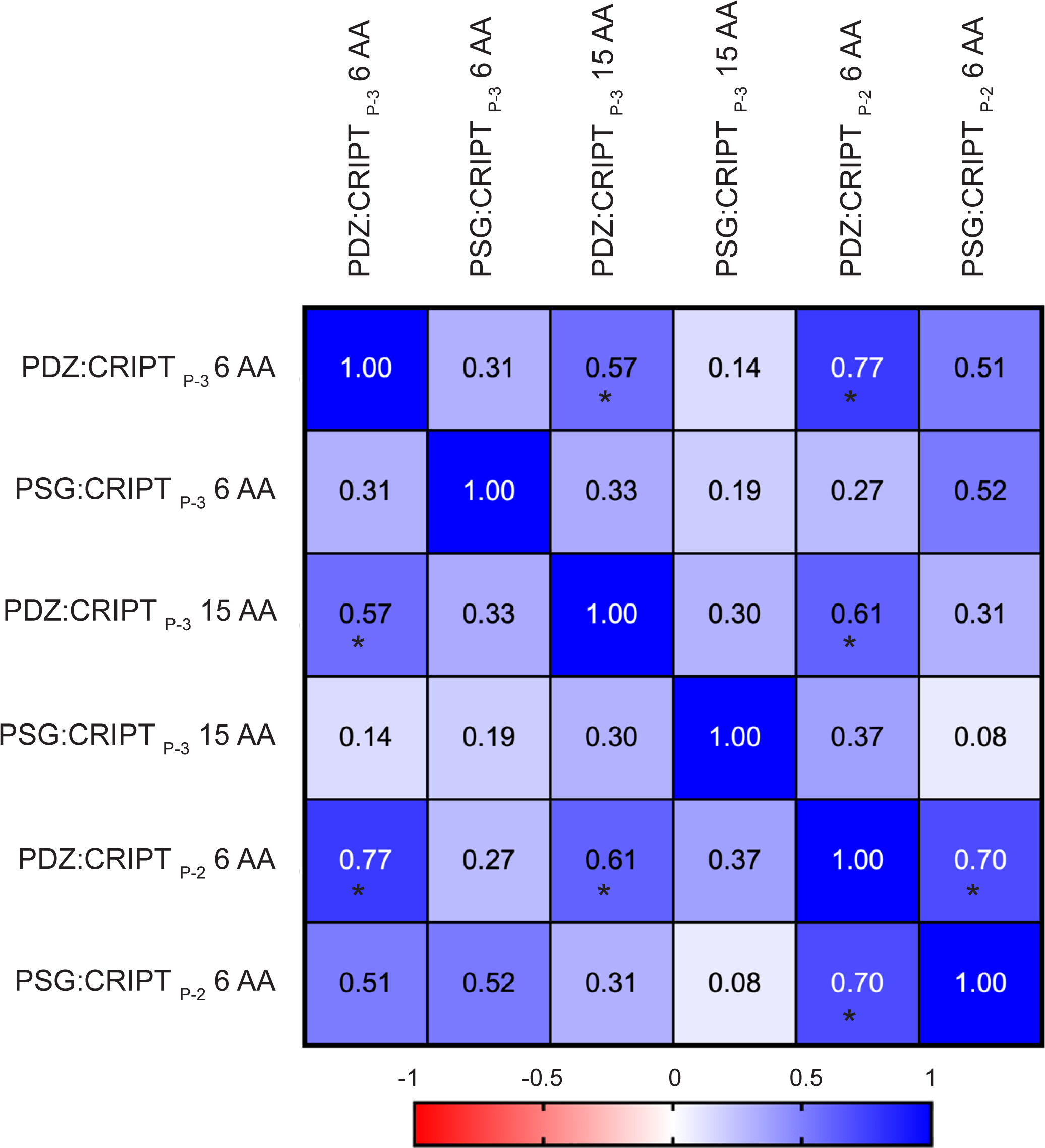
Spearman correlation analysis of the six allosteric networks involving PDZ3. The diagram reports Spearman rank correlation values for six allosteric networks in single domain PDZ3 or PSG for 14 single mutations in PDZ3 and the three peptides CRIPT_P-2_ 6 AA, CRIPT_P-3_ 6 AA or CRIPT_P-3_ 15 AA. The colour code going from dark blue (positive) to red (negative) shows the direction and strength of any monotonic relationship. A significant correlation is marked by *.

Finally, we applied the Spearman correlation analysis to the 6 allosteric networks obtained from PDZ3:CRIPT_P-2_ 6 AA, PDZ3:CRIPT_P-3_ 6 AA, PDZ3:CRIPT_P-3_ 15 AA, PSG:CRIPT_P-2_ 6 AA, PSG:CRIPT_P-3_ 6 AA and PSG:CRIPT_P-3_ 15 AA, and including 14 point mutants of PDZ3 (excluding F400A). Four significant Spearman correlations are reported, where PDZ3:CRIPT_P-2_ 6 AA are part of three monotonic relationships (Figure 4). This finding is not unexpected, as the PDZ3:CRIPT_P-2_ 6 AA allosteric network has four positions displaying positive coupling free energies. On the other hand, the PSG:CRIPT_P-3_ 6 AA allosteric network has only negative coupling free energies and while a trend is present between the PSG:CRIPT_P-3_ 6 AA and PSG:CRIPT_P-2_ 6 AA networks, there is no significant monotonic relationship.

### Correlation between coupling free energy and distance is different for PDZ3 and PSG

Another striking difference between our data sets is the distance dependence of ΔΔΔG_c_ values. In the trivial case with a non-allosteric globular protein we would expect coupling free energies to decrease with distance between the two mutations. For example, two directly interacting residues (distance <5 Å) will couple whereas residues separated by 20 Å will not. The coupling free energy between residue pairs in the PDZ3 and PSG complexes was plotted against C_α_-C_α_ distance (Figure 5). For PDZ3:CRIPT a rather clear trend was observed in which residues with high coupling free energy were located closer to the binding pocket, both for the CRIPT 6 AA and CRIPT 15 AA (Figure 5A, B and E). On the other hand, for PSG:CRIPT 6 AA we observed a peculiar inverse trend where proximal residues display ΔΔΔG_c_ values close to zero or slightly positive whereas distal residues displayed larger and negative ΔΔΔG_c_ values (Figure 5C and D). For PSG:CRIPT 15 AA the magnitude of ΔΔΔG_c_ values appeared independent of distance. In a previous study with PDZ3:CRIPT 6 AA and two other peptide mutations (P_-2_ Thr→Ser and P_0_ Val→Abu), no clear trend was observed between coupling free energy and distance (22). Altogether, the distance dependence of coupling free energies in PDZ3:CRIPT complexes are context dependent, both with regard to adjacent domains, length of peptide ligand and the precise mutation in the peptide. Overall, this variation supports our earlier notion that sequence rather than topology determines allosteric networks as quantified by coupling free energies (22, 25, 37). The noticeable change in coupling free energies from positive in PDZ3 to mainly negative in PSG underscores the hypothesis that residues rather than backbone transfer the coupling free energy (25). PSG CRIPT_P-2_ 6 AA reports positive coupling free energies for residues located next to the perturbation site, whereas the rest of the residues have negative ΔΔΔG_c_ values. Strikingly, the PSG:CRIPT_P-3_ 6 AA interaction only reports negative and neutral ΔΔΔG_c_ values. Negatively coupled residues are primarily located at the intra-domain interface between PDZ3 and SH3 domain in the supramodule PSG (38). Naganathan et al. showed, in a re-analysis of 49 structural perturbation data sets in 28 different proteins, a universal pattern in which the effect of a perturbation decays exponentially with distance from the perurbation (39). Therefore, we subjected our data set to the same analysis and plotted the numerical (absolute) value of ΔΔΔG_c_ against distance for all six data sets and fitted to an exponential function (Figure 5G). No clear trend was observed between distance and magnitude of perturbation (|ΔΔΔG_c_|) from our double mutant cycle analysis, consistent with a sequence-dependent allosteric behavior.

**Figure 5:**
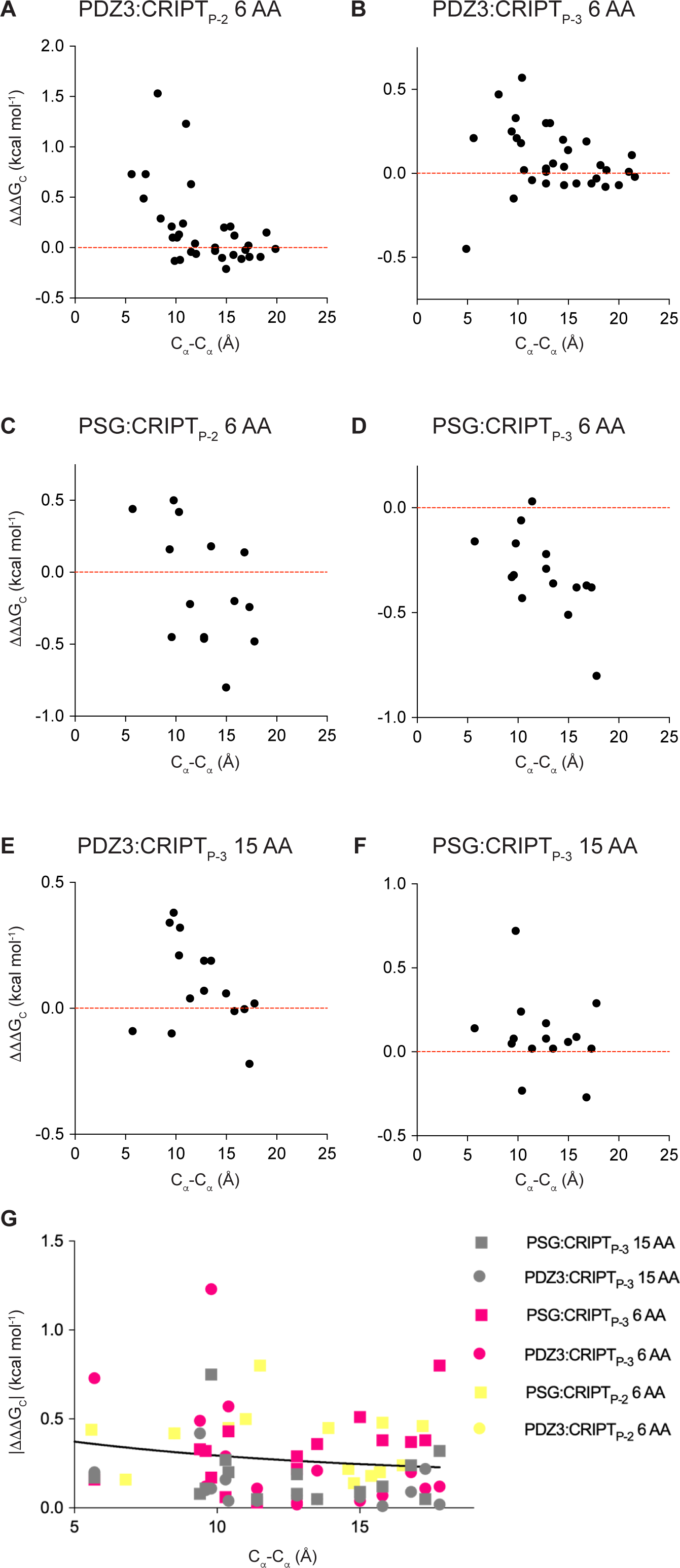
Distance dependence of coupling free energies between CRIPT and PDZ3 or PSG. The coupling free energies were plotted versus C_α_C_α_ distance between the mutation in PDZ3 and CRIPT, respectively, for the six different allosteric networks: A) PDZ3:CRIPT_P-2_ 6 AA, B) PDZ3:CRIPT_P-3_ 6 AA, C) PSG:CRIPT_P-2_ 6 AA, D) PSG:CRIPT_P-3_ 6 AA, E) PDZ3:CRIPT_P-3_ 15 AA and F) PSG:CRIPT_P-3_ 15 AA. The red horizontal line divides negative from positive coupling free energies. G) Exponential fit showing no significant distance dependence for |ΔΔΔG_c_| values from our double mutant cycles in PDZ3 and PSG (<d_c_> = 27 ± 20 Å).

## Discussion

Whilst it is well established that many biological processes are finely regulated by the allosteric nature of protein domains, a quantitative description of the energetic communication between two distinct sites in a protein is still very challenging. Nevertheless, unveiling the subset of residues that form a physically connected network between two distinct sites can have multiple applications, e.g., for protein engineering or finding a more effective and selective drug discovery approach, as allosteric sites are less conserved than the primary binding pocket within in a given protein family (40). In the case of PDZ domains, it has been shown how they represent interesting drug targets in a range of diseases including stroke and cancer. But there is no approved drug for a PDZ domain due to their promiscuity, which is manifested as binding to several similar ligands through C-terminal recognition. The presence of an evolutionarily conserved sparse allosteric network in PDZ3 was proposed two decades ago based on multiple sequence alignments and statistical coupling analysis, showing an inter residue communication from the binding pocket to a proposed allosteric site located around β_1_ and β_2_ (8). However, there is still limited application of the allosteric network in PDZ engineering and drug design as the presence of one conserved allosteric network in the PDZ family was challenged by various in silico and experimental approaches. In fact, the growing number of distinct allosteric networks in PDZ3 suggested that the choice of method influences the identity of the network (9).

### PDZ domains studied in isolation

PDZ domains play a role in protein targeting and protein complex assembly and are often found in multi domain proteins. Despite this, most *in vitro* studies use single PDZ domains to reduce complexity. Moreover, inspection of the subset of allosteric networks for PDZ3 reported in Supplementary Figure 1 shows that eight different studies have used different construct length of the PDZ3 domain, which may also affect the mapped allosteric network. One example is the α_3_ helix, which was only included in six constructs, although it is important for both binding affinity (15), folding (28) and dynamics (29). PDZ3 shares overall fold with other PDZ domains, but the α_3_ helix is exclusive for PDZ3 domains from MAGUK proteins. Thus, methods that rely on a multiple sequence alignment of all PDZ domains cannot include the α_3_ helix, which might be a limitation.

### Supertertiary structure affects the allosteric network in PDZ3

In the present study, we addressed the role of supertertiary structure by mapping the allosteric network in isolated PDZ3 as well as in the PSG supramodule. Surprisingly, we observe that not only α_3_ affects the allosteric network in PDZ3, but there is a substantial difference when the domain is studied in isolation as compared to when it is part of the PSG supramodule. This finding is in line with previous studies indicating that the allostery is mediated through interdomain communication by protein-protein interactions (15, 18).

To visualize how the presence of the SH3 and GK domains affect the allosteric network in PDZ3 (Figure 1C), we calculated ΔΔΔΔG_c_ values by subtracting ΔΔΔG_c_ for PDZ3 from ΔΔΔG_c_ for PSG (Figure 6). ΔΔΔΔG_c_ values were mapped onto the structure of PDZ3 for CRIPT_P-3_ 6 AA, CRIPT_P-2_ 6 AA and CRIPT_P-3_ 15 AA. The allosteric networks appear different, especially at the α_3_ helix, and β_1_β_2_ and β_2_β_3_ loops, as represented by the dark blue color (Figure 6A and B). In accordance with previous findings (18, 38) we propose that the allosteric network propagates to new sites in the context of the supramodule. Zhang et al. showed that the PSG from PSD-95 changes from compact to extended conformation upon CRIPT binding, which disturbs PDZ3:SH3 Interdomain interactions at the β_1_β_2_ loop, β_2_ sheet, α_2_ and α_3_ helix in PDZ3 (38). The interdomain dynamics of PDZ3:SH3 indicates that the PSG supramodule is flexible and changes interdomain conformation in response to ligand binding (38), which is supported by our results and other recent findings showing that intramolecular domain dynamics regulate the binding properties and conformations of the PSG supramodule (18). However, the interactions mediated by the β_1_β_2_ and β_2_β_3_ loops could be a general feature for PDZ supertertiary organization as reported for the PDZ1-2 tandem (41). Taken together, the data show the importance of three structural elements for inter- and intradomain structural conformation and communication: the α_3_ helix, and the β_1_β_2_ and β_2_β_3_ loops.

**Figure 6:**
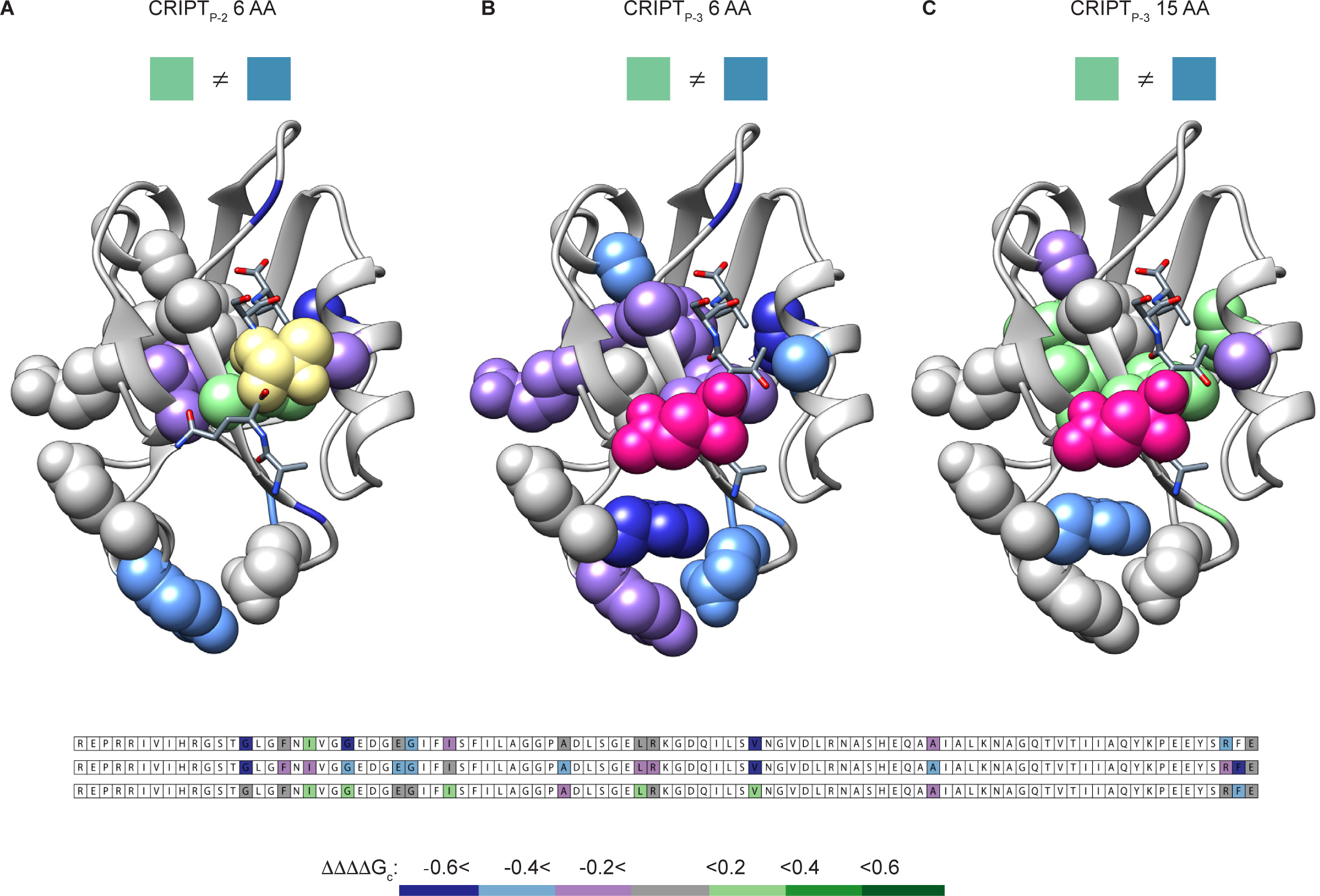
Differences between the allosteric networks in PDZ3 and PSG. The coupling free energies obtained with the single PDZ3 domain were subtracted from those of PSG for the corresponding positions to visualize the effect of SH3-GK on the allosteric network propagated in PDZ3 upon CRIPT binding. The three panels show the resulting ΔΔΔΔG_c_ values for the respective CRIPT peptide and for each tested residue, mapped onto the PDZ3:CRIPT complex. A) CRIPT_P-2_ 6 AA, B) CRIPT_P-3_ 6 AA and C) CRIPT_P-3_ 15 AA. Color code for ΔΔΔΔG_c_ values: dark blue (< −0.6), light blue (< −0.4), purple (< −0.2), grey (−0.2 to 0.2), light green (>0.2), medium green (> 0.4) and dark green (>0.6). Thus, residues with grey color show a similar response in PDZ3 and PSG to the mutational perturbation of the allosteric network.

### The significance of positive ΔΔΔG_c_ values for allostery and selectivity

Selectivity of single domains can be affected by long range allosteric networks as shown for SH3 (42, 43) as well as PDZ; e.g., PDZ1 from PTP-BL modulates the binding of ligands to the adjacent PDZ2 (42). We have previously argued that positive ΔΔΔG_c_ values are a signature for optimization of binding a certain ligand (22). Such a trend was reported here for the allosteric networks with CRIPT_P-2_ 6 AA, CRIPT_P-3_ 6 AA, and CRIPT_P-3_ 15 AA in PDZ3. Thus, a positive sign of ΔΔΔG_c_ means that a mutation in PDZ3 will result in a smaller effect of the second mutation (in CRIPT) (Figure 1B). In other words, mutation disrupts an optimized interaction network and relaxes the system with regard to further perturbation. In contrast to the mainly positive ΔΔΔG_c_ values for PDZ3, surprisingly, we observe primarily negative ΔΔΔG_c_ values for the allosteric network in PSG probed by CRIPT_P-3_ 6 AA, suggesting a non-optimized system. A tendency for negative ΔΔΔG_c_ values were recently reported for an SH3 domain (34). Thus, the presence of allosteric networks may not necessarily be linked with binding selectivity but may also trace the communication pathways between energetically interacting domains in supramodules. Analogously, the negative values in PDZ3 indicate a complex role for such domains with multiple binding partners in the presence of the α_3_ extension or the SH3-GK domains (15, 44). We note in this context that a more complex supertertiary structure may have several native, or non-native states, which would affect the allosteric coupling (45).

### Concluding remarks: what is next for allosteric networks in PDZ3?

Is there one unique or multiple allosteric networks in PDZ3? Allosteric networks have been mapped in PDZ3 using different primary structures as input and analyzed by various methods (Supplementary Figure 1), which has hampered a direct comparison. We demonstrate unequivocally that the allosteric network as defined by ΔΔΔG_c_ values is different for PDZ3 present as a single domain and in a supertertiary structure, the PSG supramodule. Therefore, we encourage using multidomain constructs in experimental and theoretical studies on intradomain allosteric networks, in particular when the domain is present in a supertertiary structure, like PDZ3. Transiting from single to multiple domains involves experimental challenges and previous focus on single domains may have hampered drug discovery. Nevertheless, the complexity and dynamics of multiple domains can offer new opportunities to control protein-protein interactions (46) and hopefully escalate attempts to profit from allosteric networks in PDZ drug design.

## Methods

### Protein expression and Purification

Wild-type PSD-95 PDZ3-SH3-GK (PSG), PDZ3 domain and mutants (all pseudo, engineered F337W in PDZ3 domains) are encoded in modified pRSET vector (Invitrogen) and transformed in Escherihia coli BL21(DE3) pLys cells (Invitrogen). Cells were grown in LB medium at 37°C and over-expression of protein was induced with 1 mM IPTG overnight at 18°C. Cells were harvested at 4°C and pellet re-solubilized in 50 mM Tris pH 7.8, 10 % glycerol (and including 100 mM NaCl for PSG) and stored at −20°C. Pellets were sonicated 2 × 4 min followed by centrifugation, 50,000 *g* at 4°C for 1 h. Supernatants were filtered and added to pre-equilibrated (50 mM Tris, 10% glycerol (including 100 mM NaCl and 0.5 mM DTT for PSG)) Nickel Sepharose Fast Flow column (GE Healthcare). Proteins were eluted with 250 mM imidazole and dialysed overnight into 50 mM Tris pH 7.8 (PDZ3) or 50 mM Tris, 2 mM DTT, 100 mM NaCl, 10% glycerol (PSG). PDZ3 or mutants were loaded onto a QS sepharose column for further purification and eluted with a 500 mM NaCl gradient. PSG or mutants were concentrated and further purified using size exclusion chromatography (S-100, GE Healthcare). Protein purity was quantified by SDS-PAGE and identity by MALDI-TOF mass spectrometry. Concentration of proteins were estimated from the absorbance at 280 nm. Circular dichroism (CD) scans of PDZ3, PSG and mutants were performed to determine if all proteins were folded. All CD experiments were performed in 50 mM sodium phosphate pH 7.45, 21 mM KCl (I = 150) at 10°C on a JASCO 1500 spectropolarimeter using the average of 5 scans. Far-UV spectra were recorded from 260 nm to 200 nm with 10 μM protein.

### Peptide synthesis

Dansyl labelled CRIPT peptides were ordered (Zhejiang Ontores Biotechnologies) or synthesized manually by SPPS as described previously (33). Concentrations were measured at 340 nm using an extinction coefficient of 3400 M^-1^ cm^-1^ (47). Based on carbon-chain length but without functional groups (OH and amide) unnatural amino acids were introduced at position P_-2_: Thr→Abu (Aminobutyric acid, R = CH_2_CH_3_) and positionP_-3_: Gln→Nva (Norvaline, R = CH_2_CH_2_CH_3_).

### Kinetic experiments

All kinetic experiments were performed in 50 mM sodium phosphate pH 7.45, 21 mM KCl (I = 150) at 10°C (including 0.5 mM Tris-2-carboxyethyl-phosphine (TCEP) for experiments with PSG). Binding and dissociation experiments were performed using an upgraded SX-17 MV stopped-flow spectrophotometer (Applied Photophysics, Leatherhead, UK) and carried out as described previously (48).

Association rate constants (*k*_on_) were obtained from binding experiments under conditions approaching pseudo first order at high CRIPT concentrations (dansylated) ([PDZ or PSG]<[CRIPT]). Thus, PDZ (1μM) were mixed rapidly with increasing concentrations of dansylated CRIPT ligand (2 to 20 μM of CRIPT 6 AA, CRIPT_P-2_ 6 AA, CRIPT_P-3_ 6 AA, CRIPT 15 AA, CRIPT_P-2_ 15 AA or CRIPT_P-3_ 6 AA). Excitation was at 280 nm and emission measured around 330 nm (using a 330 ± 25 nm interference filter). Kinetic traces (minimum 5) were recorded, averaged, and fitted to a single exponential function (a double exponential was used for PSG with CRIPT 6AA, CRIPT_P-2_ 6 AA and CRIPT_P-3_ 6 AA) consistent with a two-state bimolecular association/dissociation mechanism. The fitting yielded observed rate constants (*k*_*obs*_) for each CRIPT concentration. A plot with *k*_*obs*_ versus CRIPT concentration was obtained and fitted to an equation derived for a second order bimolecular association reaction to account for the deviation from linearity at low CRIPT concentrations.

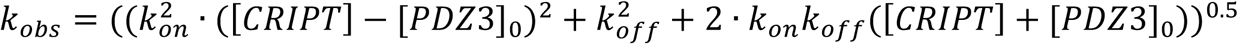

[PDZ3]_0_ and [CRIPT] are the initial protein and peptide concentrations. While the association rate constant (*k*_*on*_) was obtained from fitting of this equation, the dissociation rate constant (*k*_*off*_) was obtained from displacement experiments. PDZ or PSG (2 μM) was mixed with dansylated CRIPT (10 μM) followed by a long incubation (>15 min) to ensure that equilibrium was established. Dansylated CRIPT was displaced from PDZ by mixing with an excess of unlabelled CRIPT (100, 150, 200 μM). *k*_*off*_ values were estimated from the average of three *k*_*obs*_ determinations at high concentrations of unlabelled CRIPT peptide, where the *k*_*obs*_ values were constant with unlabelled CRIPT concentration. All binding and displacement experiments performed with PSG and CRIPT 6 AA were fitted to double exponential function due to more complex binding mechanism, and *k*_*obs*_ values from the fast phase in the respective experiment were used for further analysis (30).

### Isothermal Titration Calorimetry

All ITC experiments were performed in 50 mM sodium phosphate pH 7.45, 21 mM KCl (I = 150 mM) at 25°C, and including 0.5 mM TCEP for experiments with PSG, in an iTC200 instrument (Malvern). CRIPT and PDZ or PSG were dialysed overnight in the same buffer to minimize artifacts from buffer mismatch in the experiment. CRIPT was titrated (16-19 injections) into a cell with PDZ or PSG, and saturation was obtained by having 10-20 higher concentration of CRIPT in the syringe resulting in a roughly 2-fold excess of CRIPT at the end of the titration. Baseline corrections were performed to minimize the chi-value in curve fitting. Fitting of data was performed using the iTC200 software.

### Calculation of coupling free energies

Coupling free energies were calculated from *K*_*d*_ values determined by the ratio *k*_*off*_/*k*_*on*_ for all four complexes in the double mutant cycle (Figure 1B) as described (21, 22). Concisely, four complexes are characterized for each double mutant cycle: 1 (WT), 2 (single mutation in PDZ or PSG), 3 (single mutation in CRIPT) and 4 (single mutation in PDZ/PSG and CRIPT). The change in free energy upon single point mutation in PDZ, can be measured by calculating the difference in energy from complex 1 to 2 (Eq. 1). Similarly, the energy change upon single point mutation in PDZ bound to mutated CRIPT, can be measured by calculating the difference in energy from complex 3 to 4 (Eq. 2). Therefore, if the residue subjected to single point mutation in PDZ does not interact energetically with the residue mutated in CRIPT, the energy difference upon single point mutation in PDZ will be similar for the two complexes 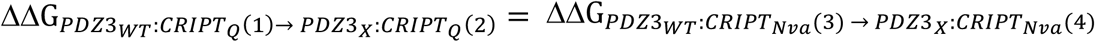 and meaning that the coupling free energy (ΔΔΔG_c_) is zero (Eq. 3). However, if the residues subjected to single point mutations in PDZ (X) and CRIPT (Nva) interact energetically the calculated ΔΔΔG_c_ ≠ 0. This coupling free energy is then a quantitative measure of the strength of the interaction between the two residues, one in PDZ and one in CRIPT.

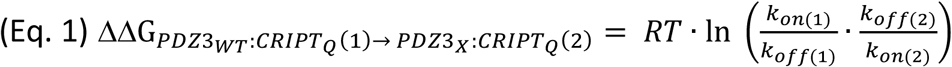

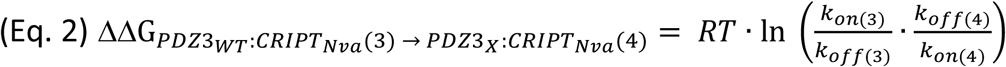

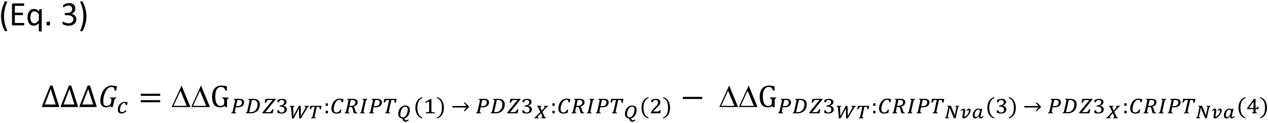

## Acknowledgements

The project has received funding from the European Union’s Horizon 2020 research and innovation programme under the Marie Sklodowska-Curie grant agreement No 67341 (to SG and PJ) and from the Swedish Research Council (grant no. 2016-04965) (to PJ).

## Supplementary material

**Supplementary Figure 1.**
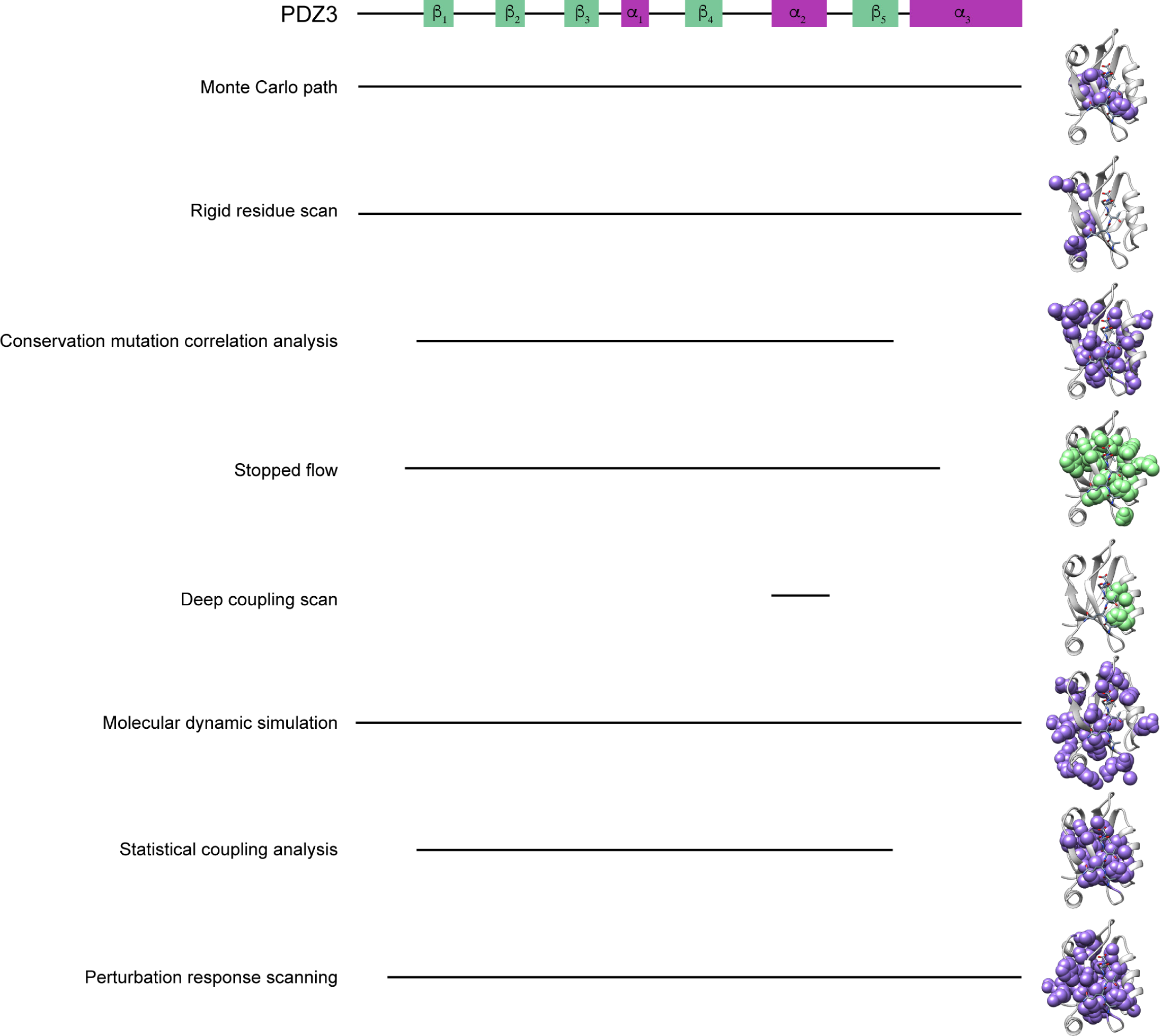
Allosteric networks in PDZ3 determined by various methods. Overview showing 8 different allosteric networks reported in PDZ3 using in silico approaches (six examples, purple) and two experimental approaches (green). The illustration highlights the different lengths of PDZ3 constructs used for the different approaches: Monte Carlo path [1], rigid residue scan [2], conservation mutation correlation analysis [3], thermodynamic double mutant cycle by stopped flow [4], deep coupling scan [5], molecular dynamics scan [6], statistical coupling analysis [7], perturbation response scanning [8]. The network illustrations were adapted from [9].

**Supplementary Figure 2.**
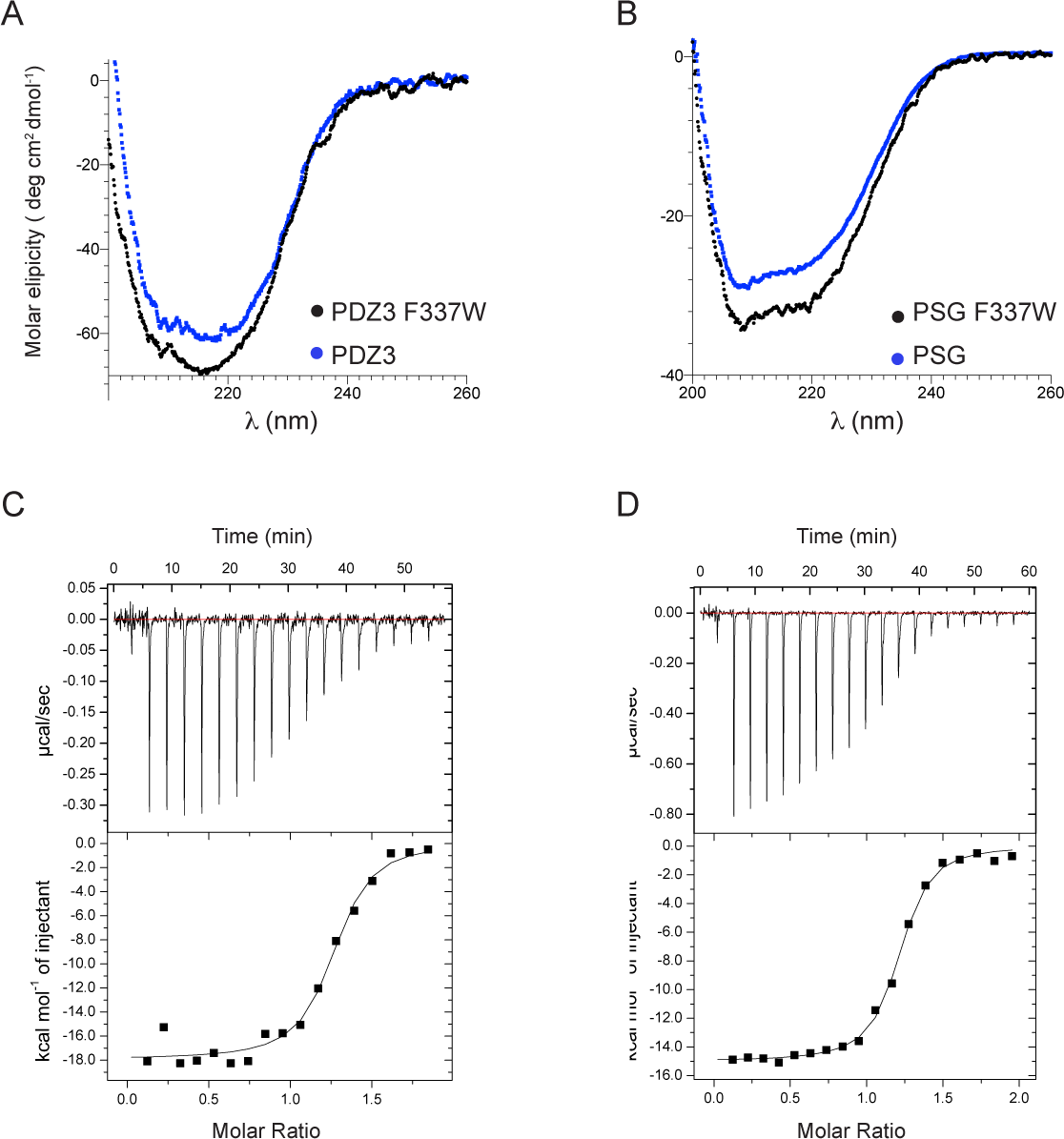
F337W in PDZ3 and PSG does not affect secondary structure and affinity of CRIPT. Circular dichroism spectra at 200-260 nm for PDZ3 (A) and PSG (B) were recorded in 50 mM sodium phosphate pH 7.4, 10°C. ITC experiments for CRIPT 6 AA binding to (C) PSG and (D) PSG with F337W probe were performed in 50 mM sodium phosphate pH 7.4, 25°C.

**Supplementary Figure 3.**
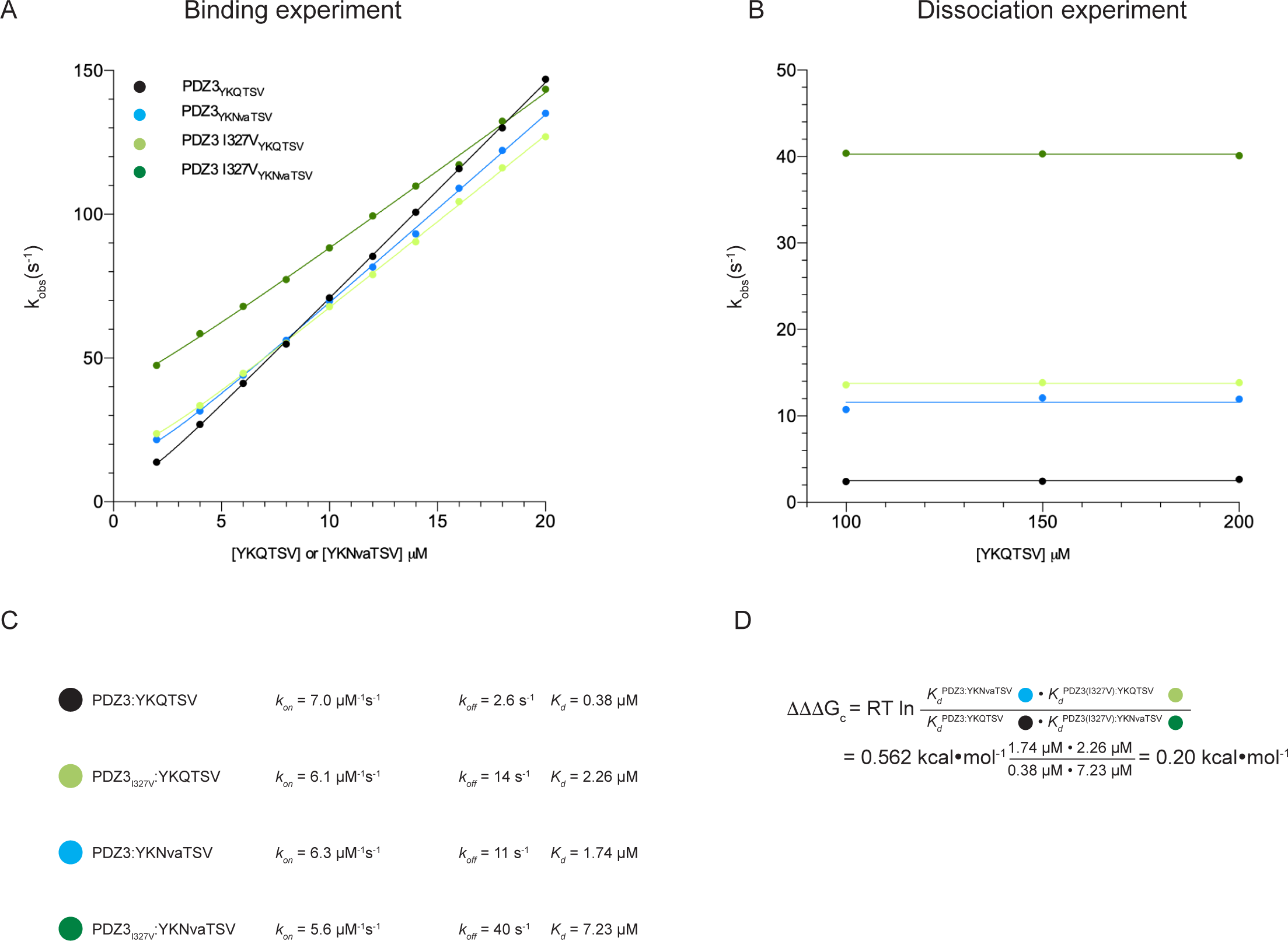
Calculation example of coupling free energy. Calculation example of coupling free energy (ΔΔΔG_c_) from observed rate constants for WT and I327V PDZ3. A) Plot showing the linear relationship between observed rate constants and CRIPT 6 AA concentration under pseudo first order conditions. The association rate constant (*k*_*on*_) is equal to the slope of the curve. The dissociation rate constant (*k*_*off*_) was estimated from a separate displacement experiment. B) Plot showing observed rate constants from displacement experiments at three concentrations of unlabelled CRIPT 6 AA. Displacement experiments were performed by preincubating PDZ3:CRIPT 6 AA (dansylated) and mixing with 100, 150, or 200 μM unlabelled CRIPT 6 AA. At high concentration the dissociation of labelled CRIPT 6 AA is practically irreversible and the average of three observed rate constants was used as an estimate of the dissociation rate constant. C) The *K*_*d*_ value was calculated from the ratio of the dissociation and association rate constants for the four complexes. D) Calculation example of coupling free energy (ΔΔΔG_c_) for the double mutant cycle probing Gln_-3_ in CRIPT 6 AA and I327 in PDZ3.

**Supplementary Figure 4.**
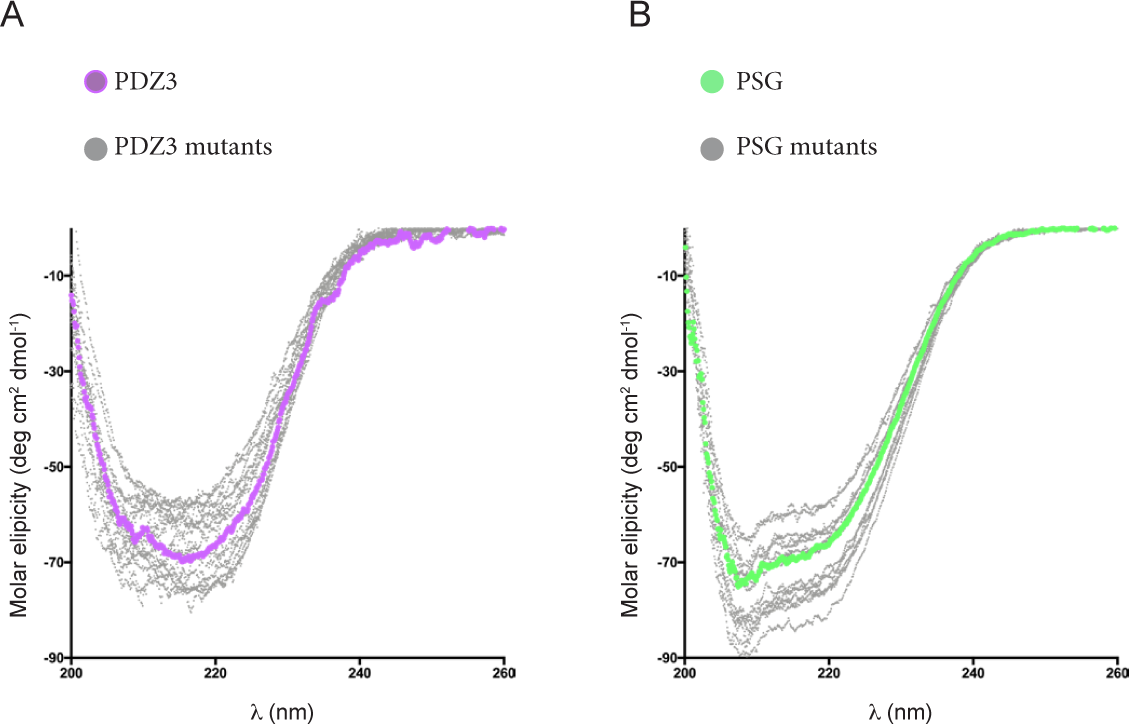
The Mutations did not perturb secondary structure of PDZ3 and PSG. CD spectra shown for 15 mutants of PDZ3 (A) and PSG (B). Color code: PDZ3 WT purple, PSG WT green and mutants grey. The spectra were recorded in 50 mM Sodium Phosphate pH 7.4, 10°C including 0.5 mM TCEP for PSG.

**Supplementary Figure 5.**
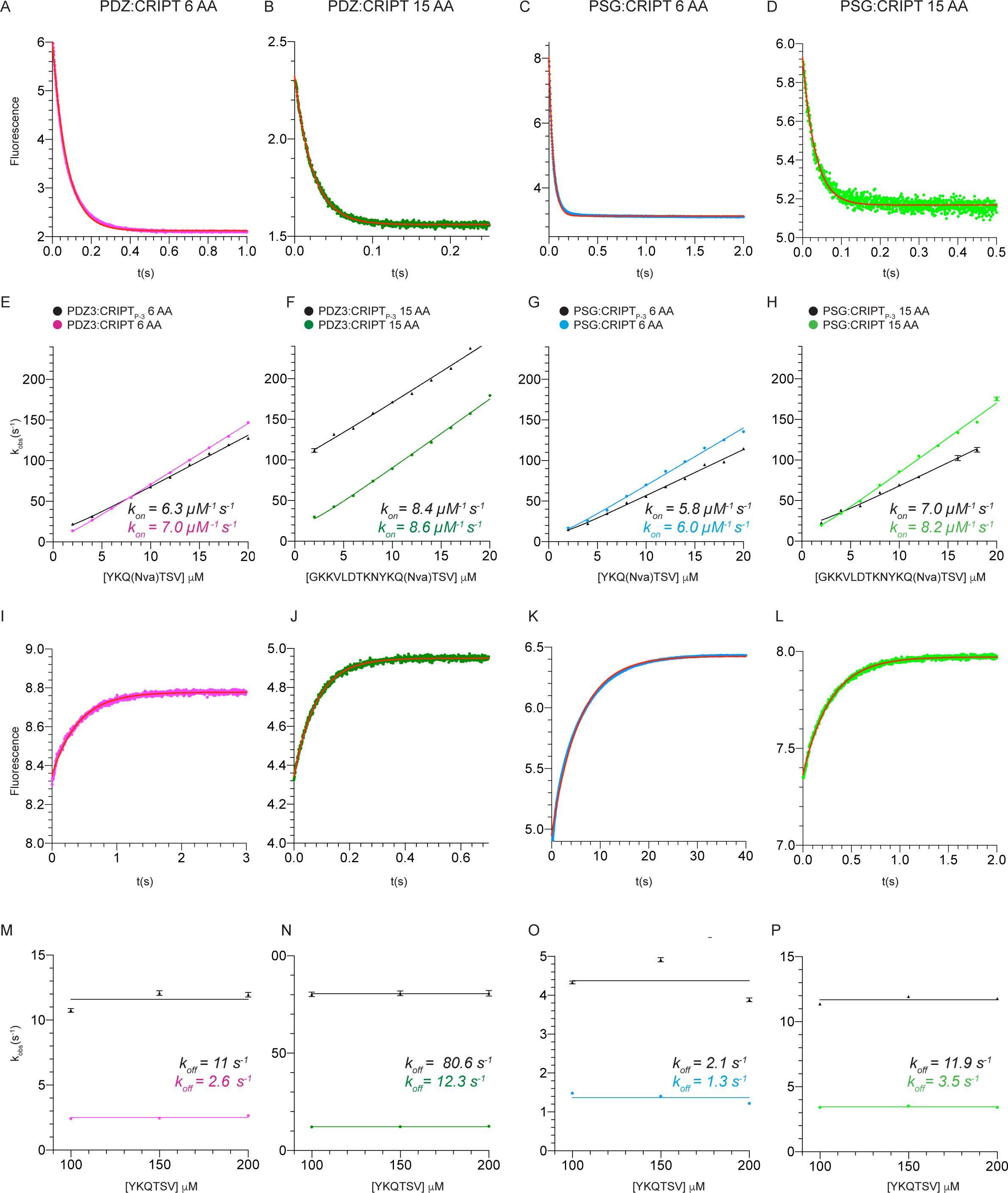
Binding and dissociation kinetics of CRIPT peptides and PDZ3 or PSG. A-D) Examples of kinetic traces from stopped flow experiments for binding of dansylated CRIPT (4μM): CRIPT 6 AA (A and C) or CRIPT 15 AA (B and D) to PDZ3 (1 μM) (A and B) or PSG (1 μM) (C or D). Traces were fitted to single (A, B and D) or double exponential functions (C). E-H) The experiments were repeated over a range of dansylated CRIPT concentrations (2 to 20 μM) at a constant concentration of PDZ3 or PSG. The observed rate constant (*k*_*obs*_) was obtained from the fit to single or double exponential functions and plotted versus [CRIPT]. From the linear relationship between [CRIPT] and *k*_*obs*_ the associations constant (*k*_*on*_) was obtained as the slope rate. I-L) Examples of kinetic traces from stopped flow experiments in which *k*_*off*_ was determined in a displacement reaction. A large excess of unlabeled CRIPT was mixed with pre-equilibried pro-tein:peptide complex (2:10 µM) of the respective PDZ3 or PSG and dansyl labelled CRIPT. Traces were fitted to a single (I, J and L) or double (K) exponential function to obtain the observed rate constant. M-P) The experiments were repeated with three different concentrations of unlabeled CRIPT (100, 150 and 200 μM) and an average of the observed rate constants was used as the dissociation rate constant (*k*_*off*_). Color codes for protein:peptide complex: pink (PDZ3:CRIPT 6 AA), green (PDZ3: CRIPT 15 AA), blue (PSG:CRIPT 6 AA), light green (PSG:CRIPT 15 AA) and black (the same peptide but with P_-3_ (Nva) mutation). All fitted parameters are presented in Supplementary Tables 2-3 and 5-6.

**Supplementary Figure 6.**
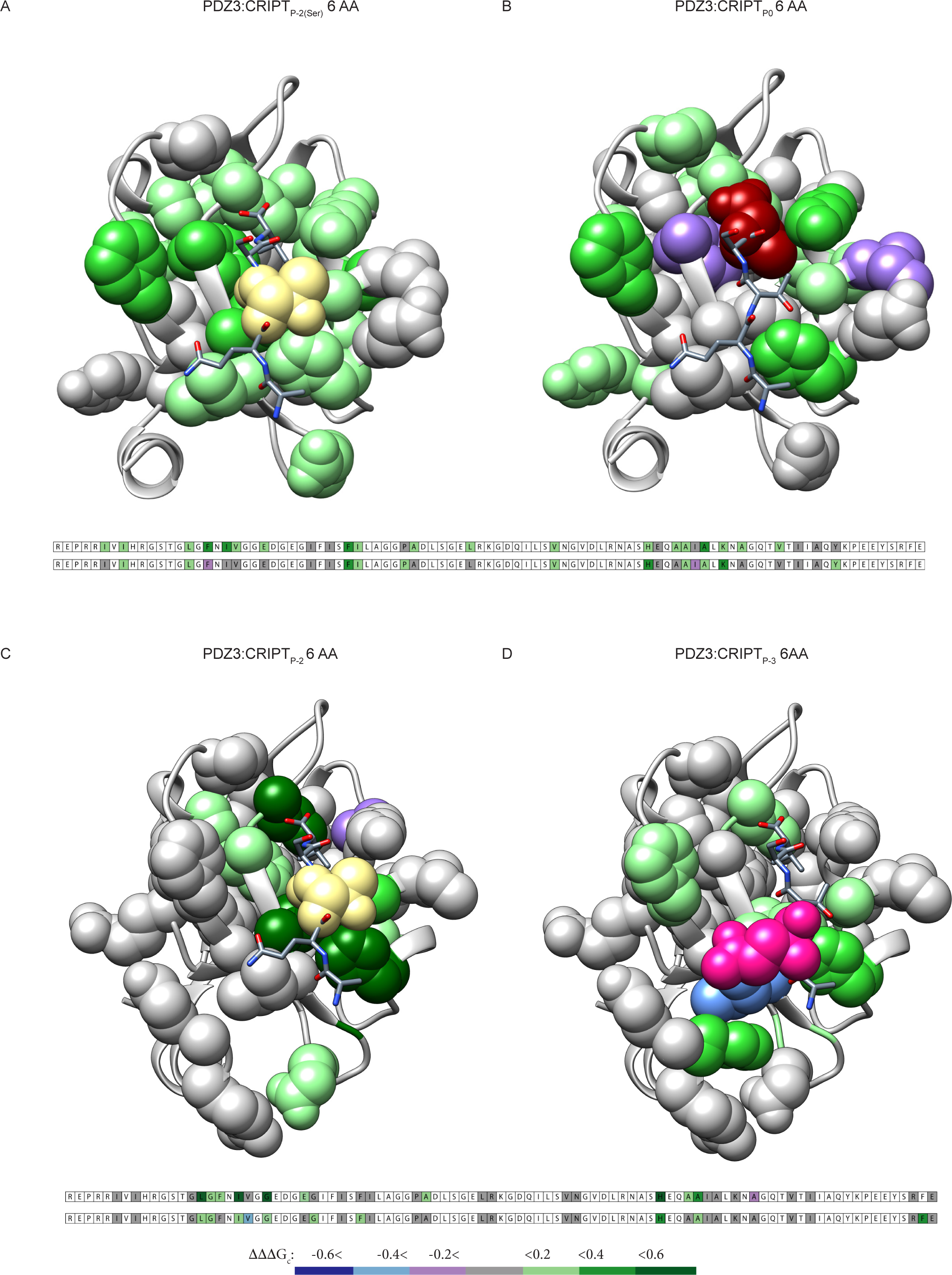
Origin of perturbations affect the allosteric network in PDZ3. Comparison of mapped allosteric networks (ΔΔΔG_c_) in PDZ3 for four peptide mutations at different positions: A) P_-2_: Thr to Ser B) P_0_: Val to Abu C) P_-2_: Thr to Abu and D) P_-3_: Gln to Nva. The mapped allosteric networks reported in A and B were published previously [4], whereas C and D is the present work.

**Supplementary Figure 7.**
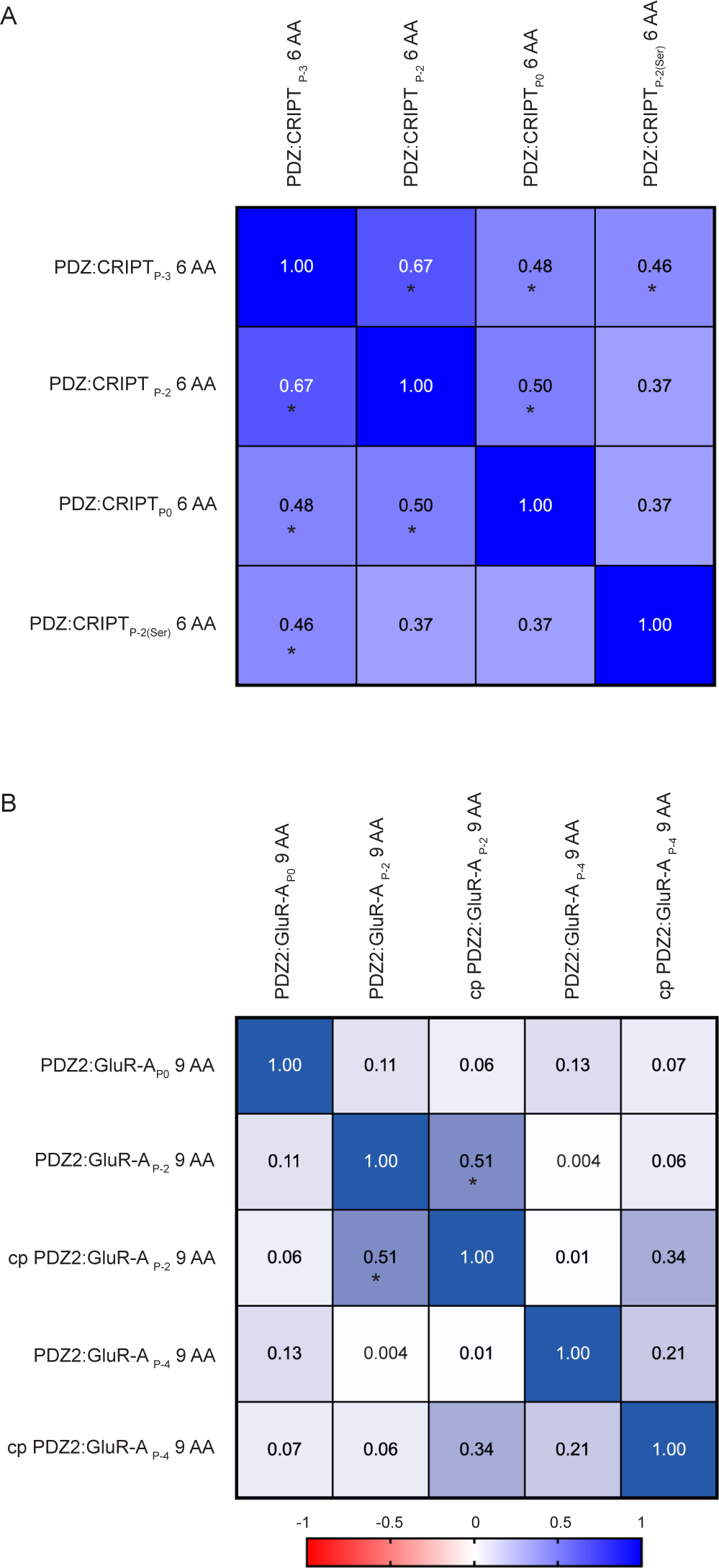
Spearman correlation analysis of allosteric networks involving PDZ3. The diagram reports Spearman rank correlation values for A) four allosteric networks reported in PDZ3 with 22 single mutations and four peptides: CRIPT_P0_ 6 AA [4], CRIPT_P-2_ 6 AA, CRIPT_P-2(Ser)_ 6 AA [4] and CRIPT_P-3_ 6 AA. B) five allosteric networks in SAP97 PDZ2 and a circularly permuted (cp) SAP97 PDZ2 with 19 single mutations and three peptides: GluR-A_P0_ 9 AA, GluR-A_P-2_ 9 AA and GluR-A_P-4_ 9 AA [10]. The color code going from dark blue (positive) to red (negative) shows the direction and strength of any monotonic relationship. Significant correlation coefficient marked with *.

---

**Supplementary Table 1.** Coupling free energies (ΔΔΔG_c_) for all mutated residues in PDZ3 Color code: dark blue (< −0.6), light blue (< −0.4), purple (< −0.2), grey (−0.2 to 0.2), light green (>0.2), medium green (> 0.4) and dark green (>0.6).

|  | PDZ3 | PDZ3 |
| --- | --- | --- |
|  | YKNvaTSV | YKQAbuSV |
| I314V | - 0.02 | - 0.01 |
| I316A | 0.01 | - 0.09 |
| G322A | 0.14 | - 0.04 |
| L323A | 0.20 | 0.63 |
| G324A | 0.30 | 0.24 |
| F325A | 0.18 | 0.29 |
| I327V | 0.21 | 0.73 |
| V328A | - 0.45 | 0.10 |
| G330A | 0.33 | 1.23 |
| E334A | 0.06 | 0.21 |
| G335A | 0.30 | - 0.03 |
| I338A | - 0.15 | - 0.12 |
| F340A | 0.21 | 0.10 |
| I341V | 0.03 | 0.04 |
| P346G | 0.02 | 0.003 |
| A347G | 0.19 | 0.20 |
| L353A | - 0.06 | - 0.11 |
| R354A | - 0.06 | - 0.07 |
| V362A | - 0.03 | 0.12 |
| N363A | 0.11 | 0.15 |
| H372A | 0.47 | 1.53 |
| A375G | 0.02 | 0.21 |
| A376G | 0.25 | 0.49 |
| I377A | 0.01 | - 0.13 |
| A378G | - 0.07 | - 0.06 |
| K380A | 0.04 | 0.13 |
| A382G | 0.05 | - 0.21 |
| V386A | - 0.07 | - 0.09 |
| I388V | - 0.08 | - 0.02 |
| R399A | - 0.06 | 0.02 |
| F400A | 0.57 | N.D. |
| E401A | - 0.04 | - 0.10 |

**Supplementary Table 2.** *k*_*on*_, *k*_*off*_ and *K*_*d*_ for the interaction between PDZ3 and CRIPT 15 AA or CRIPT _P-3_ 15 AA.

|  | GKKVLDTKNYKQTSV |  |  | GKKVLDTKNYKNvaTSV |  |  |
| --- | --- | --- | --- | --- | --- | --- |
| | $k_{on}$ (s<sup>-1</sup> μM<sup>-1</sup>) | $k_{off}$ (s<sup>-1</sup>) | $K_d$ (μM) | $k_{on}$ (s<sup>-1</sup> μM<sup>-1</sup>) | $k_{off}$ (s<sup>-1</sup>) | $K_d$ (μM) |
| PDZ3 | 8.6 ± 0.1 | 12 ± 0.1 | 1.4 ± 0.03 | 8.4 ± 0.2 | 81 ± 0.2 | 9.7 ± 0.3 |
| G322A | 8.0 ± 0.1 | 3.5 ± 0.02 | 0.4 ± 0.01 | 7.6 ± 0.1 | 20 ± 0.1 | 2.7 ± 0.03 |
| F325A | 11 ± 0.2 | 26 ± 0.2 | 2.4 ± 0.1 | 11 ± 0.6 | 125 ± 6 | 11 ± 0.8 |
| I327V | 7.6 ± 0.1 | 62 ± 1 | 8.2 ± 0.2 | 6.1 ± 0.7 | 397 ± 11 | 65 ± 8 |
| G330A | 8.7 ± 0.3 | 97 ± 0.4 | 11 ± 0.4 | 8.1 ± 3.4 | 307 ± 37 | 38 ± 17 |
| E334A | 8.3 ± 0.1 | 22 ± 0.2 | 2.6 ± 0.1 | 10 ± 0.6 | 130 ± 7 | 13 ± 1 |
| G335A | 12 ± 0.2 | 4.5 ± 0.01 | 0.4 ± 0.01 | 11 ± 0.2 | 24 ± 0.3 | 2.1 ± 0.05 |
| I338A | 7.2 ± 0.1 | 11 ± 0.1 | 1.5 ± 0.03 | 5.6 ± 0.2 | 65 ± 2 | 12 ± 0.4 |
| A347G | 7.3 ± 0.2 | 25 ± 0.2 | 3.4 ± 0.1 | 5.8 ± 0.3 | 135 ± 4 | 23 ± 1 |
| L353A | 8.1 ± 0.1 | 12 ± 0.03 | 1.5 ± 0.03 | 5.8 ± 0.3 | 85 ± 1 | 15 ± 0.8 |
| R354A | 9.1 ± 0.1 | 12 ± 0.1 | 1.4 ± 0.04 | 9.4 ± 0.3 | 87 ± 2 | 9.2 ± 0.3 |
| V362A | 7.3 ± 0.1 | 9.8 ± 0.04 | 1.3 ± 0.03 | 8.3 ± 0.2 | 72 ± 2 | 8.7 ± 0.3 |
| A376G | 8.6 ± 0.2 | 39 ± 0.5 | 4.5 ± 0.2 | 11 ± 0.7 | 190 ± 8 | 17 ± 1 |
| R399A | 11 ± 0.1 | 7.8 ± 0.1 | 0.7 ± 0.02 | 11 ± 0.1 | 38 ± 0.4 | 3.4 ± 0.05 |
| F400A | 9.8 ± 0.1 | 13 ± 0.1 | 1.3 ± 0.02 | 15 ± 0.4 | 72 ± 1 | 4.9 ± 0.2 |
| E401A | 7.5 ± 0.1 | 15 ± 0.1 | 2.0 ± 0.1 | 7.8 ± 0.2 | 96 ± 0.6 | 12 ± 0.4 |

**Supplementary Table 3.** *k*_*on*_, *k*_*off*_ and *K*_*d*_ for the interaction between PDZ3 and CRIPT 6 AA or CRIPT _P-3_ 6 AA.

|  | YKQTSV |  |  | YKNvaTSV |  |  |
| --- | --- | --- | --- | --- | --- | --- |
| | $k_{on}$ (s<sup>-1</sup> μM<sup>-1</sup>) | $k_{off}$ (s<sup>-1</sup>) | $K_d$ (μM) | $k_{on}$ (s<sup>-1</sup> μM<sup>-1</sup>) | $k_{off}$ (s<sup>-1</sup>) | $K_d$ (μM) |
| PDZ3 | 7.0 ± 0.1 | 2.6 ± 0.04 | 0.4 ± 0.01 | 6.3 ± 0.1 | 11 ± 0.3 | 1.7 ± 0.1 |
| I314V | 7.1 ± 0.04 | 2.4 ± 0.04 | 0.3 ± 0.01 | 6.6 ± 0.1 | 11 ± 0.3 | 1.7 ± 0.1 |
| I316A | 6.3 ± 0.03 | 1.5 ± 0.03 | 0.2 ± 0.005 | 5.7 ± 0.1 | 6 ± 0.1 | 1.1 ± 0.02 |
| G322A | 6.2 ± 0.1 | 0.7 ± 0.02 | 0.1 ± 0.004 | 6.0 ± 0.1 | 3 ± 0.1 | 0.4 ± 0.01 |
| L323A | 7.1 ± 0.04 | 1.3 ± 0.3 | 1.8 ± 0.04 | 6.5 ± 0.1 | 37 ± 1 | 5.7 ± 0.2 |
| G324A | 6.7 ± 0.1 | 12 ± 0.3 | 1.9 ± 0.1 | 6.3 ± 0.3 | 31 ± 2 | 5.0 ± 0.4 |
| F325A | 8.0 ± 0.2 | 5.8 ± 0.2 | 0.7 ± 0.03 | 7.8 ± 0.1 | 19 ± 0.7 | 2.5 ± 0.1 |
| I327V | 6.1 ± 0.04 | 14 ± 0.1 | 2.3 ± 0.02 | 5.6 ± 0.1 | 40 ± 0.1 | 7.2 ± 0.1 |
| V328A | 7.1 ± 0.1 | 7.2 ± 0.2 | 1.0 ± 0.03 | 7.5 ± 0.1 | 79 ± 2 | 10.6 ± 0.3 |
| G330A | 6.3 ± 0.2 | 65 ± 3 | 10 ± 0.6 | 8.0 ± 0.8 | 216 ± 18 | 27 ± 3 |
| E334A | 9.2 ± 0.1 | 5.6 ± 0.1 | 0.6 ± 0.01 | 7.6 ± 0.1 | 20 ± 0.04 | 2.6 ± 0.04 |
| G335A | 14 ± 0.2 | 0.5 ± 0.03 | 0.04 ± 0.002 | 13 ± 0.2 | 1 ± 0.1 | 0.1 ± 0.005 |
| I338A | 6.0 ± 0.03 | 1.9 ± 0.04 | 0.3 ± 0.01 | 5.4 ± 0.0 | 10 ± 0.2 | 1.9 ± 0.1 |
| F340A | 6.3 ± 0.1 | 4.4 ± 0.1 | 0.7 ± 0.01 | 5.8 ± 0.1 | 13 ± 0.5 | 2.3 ± 0.1 |
| I341V | 7.0 ± 0.04 | 3.4 ± 0.1 | 0.5 ± 0.01 | 6.4 ± 0.1 | 14 ± 0.3 | 2.2 ± 0.1 |
| P346G | 6.8 ± 0.1 | 4.1 ± 0.2 | 0.6 ± 0.03 | 6.1 ± 0.1 | 17 ± 0.6 | 2.7 ± 0.1 |
| A347G | 5.8 ± 0.03 | 5.8 ± 0.3 | 1.0 ± 0.1 | 5.2 ± 0.1 | 17 ± 0.2 | 3.3 ± 0.1 |
| L353A | 6.7 ± 0.03 | 1.9 ± 0.1 | 0.3 ± 0.01 | 6.0 ± 0.1 | 9 ± 0.1 | 1.5 ± 0.03 |
| R354A | 7.9 ± 0.1 | 2.7 ± 0.04 | 0.3 ± 0.01 | 6.2 ± 0.1 | 11 ± 0.1 | 1.8 ± 0.03 |
| V362A | 5.6 ± 0.1 | 2.4 ± 0.02 | 0.4 ± 0.01 | 5.2 ± 0.1 | 11 ± 0.1 | 2.1 ± 0.04 |
| N363A | 6.8 ± 0.1 | 4.0 ± 0.1 | 0.6 ± 0.02 | 6.2 ± 0.1 | 14 ± 0.3 | 2.3 ± 0.1 |
| H372A | 5.3 ± 0.1 | 35 ± 0.4 | 6.5 ± 0.1 | 9.9 ± 0.7 | 131 ± 9 | 13 ± 1 |
| A375G | 5.1 ± 0.1 | 6.1 ± 0.1 | 1.2 ± 0.03 | 4.6 ± 0.1 | 25 ± 0.1 | 5.4 ± 0.1 |
| A376G | 7.2 ± 0.03 | 10 ± 0.4 | 1.4 ± 0.1 | 6.4 ± 0.1 | 28 ± 0.1 | 4.3 ± 0.1 |
| I377A | 6.9 ± 0.04 | 2.1 ± 0.01 | 0.3 ± 0.002 | 6.1 ± 0.1 | 9 ± 0.6 | 1.4 ± 0.1 |
| A378G | 6.5 ± 0.1 | 1.5 ± 0.01 | 0.2 ± 0.002 | 6.5 ± 0.1 | 8 ± 0.3 | 1.2 ± 0.1 |
| K380A | 5.4 ± 0.03 | 5.1 ± 0.1 | 0.9 ± 0.02 | 5.0 ± 0.1 | 21 ± 0.2 | 4.1 ± 0.1 |
| A382G | 6.9 ± 0.04 | 1.0 ± 0.01 | 0.1 ± 0.001 | 6.3 ± 0.1 | 4 ± 0.1 | 0.6 ± 0.01 |
| V386A | 6.6 ± 0.04 | 1.9 ± 0.1 | 0.3 ± 0.01 | 6.0 ± 0.0 | 9 ± 0.1 | 1.5 ± 0.02 |
| I388V | 7.2 ± 0.1 | 3.3 ± 0.04 | 0.5 ± 0.01 | 6.4 ± 0.1 | 16 ± 0.3 | 2.4 ± 0.1 |
| R399A | 8.0 ± 0.1 | 7.3 ± 0.1 | 0.9 ± 0.01 | 5.8 ± 0.1 | 27 ± 0.1 | 4.7 ± 0.1 |
| F400A | 7.5 ± 0.5 | 110 ± 4 | 15 ± 1 | 9.1 ± 4 | 229 ± 49 | 25 ± 11 |
| E401A | 7.3 ± 0.1 | 5.3 ± 0.03 | 0.7 ± 0.01 | 6.2 ± 0.2 | 22 ± 0.1 | 3.6 ± 0.1 |

**Supplementary Table 4.** *k*_*on*_, *k*_*off*_ and *K*_*d*_ for the interaction between PDZ3 and CRIPT 6 AA or CRIPT _P-2_ 6 AA.

|  | YKQTSV |  |  | YKQAbuSV |  |  |
| --- | --- | --- | --- | --- | --- | --- |
| | $k_{on}$ (s<sup>-1</sup> μM<sup>-1</sup>) | $k_{off}$ (s<sup>-1</sup>) | $K_d$ (μM) | $k_{on}$ (s<sup>-1</sup> μM<sup>-1</sup>) | $k_{off}$ (s<sup>-1</sup>) | $K_d$ (μM) |
| PDZ3 | 7.0 ± 0.1 | 2.6 ± 0.04 | 0.4 ± 0.01 | 7.5 ± 0.1 | 68 ± 0.2 | 9.0 ± 0.2 |
| I314V | 7.1 ± 0.04 | 2.4 ± 0.04 | 0.3 ± 0.01 | 7.6 ± 0.2 | 64 ± 0.6 | 8.5 ± 0.2 |
| I316A | 6.3 ± 0.03 | 1.5 ± 0.03 | 0.2 ± 0.005 | 6.3 ± 0.1 | 43 ± 0.3 | 6.8 ± 0.1 |
| G322A | 6.2 ± 0.1 | 0.7 ± 0.02 | 0.1 ± 0.004 | 6.2 ± 0.1 | 19 ± 0.2 | 3.0 ± 0.1 |
| L323A | 7.1 ± 0.04 | 1.3 ± 0.3 | 1.8 ± 0.04 | 16 ± 2 | 221 ± 26 | 14 ± 2 |
| G324A | 6.7 ± 0.1 | 12 ± 0.3 | 1.9 ± 0.1 | 6.6 ± 0.7 | 193 ± 17 | 29 ± 4 |
| F325A | 8.0 ± 0.2 | 5.8 ± 0.2 | 0.7 ± 0.03 | 9.0 ± 0.4 | 94 ± 5 | 11 ± 0.7 |
| I327V | 6.1 ± 0.04 | 14 ± 0.1 | 2.3 ± 0.02 | 12 ± 1 | 172 ± 15 | 15 ± 2 |
| V328A | 7.1 ± 0.1 | 7.2 ± 0.2 | 1.0 ± 0.03 | 8.5 ± 1 | 175 ± 9 | 21 ± 3 |
| G330A | 6.3 ± 0.2 | 65 ± 3 | 10 ± 0.6 | 7.3 ± 0.7 | 203 ± 16 | 28 ± 3 |
| E334A | 9.2 ± 0.1 | 5.6 ± 0.1 | 0.6 ± 0.01 | 9.8 ± 0.3 | 99 ± 4 | 10 ± 0.5 |
| G335A | 14 ± 0.2 | 0.5 ± 0.03 | 0.04 ± 0.002 | 15 ± 0.3 | 14 ± 0.1 | 1.0 ± 0.02 |
| I338A | 6.0 ± 0.03 | 1.9 ± 0.04 | 0.3 ± 0.01 | 6.1 ± 0.1 | 57 ± 0.3 | 9.3 ± 0.2 |
| F340A | 6.3 ± 0.1 | 4.4 ± 0.1 | 0.7 ± 0.01 | 6.7 ± 0.2 | 95 ± 1 | 14 ± 0.5 |
| I341V | 7.0 ± 0.04 | 3.4 ± 0.1 | 0.5 ± 0.01 | 8.2 ± 0.2 | 90 ± 0.4 | 11 ± 0.3 |
| P346G | 6.8 ± 0.1 | 4.1 ± 0.2 | 0.6 ± 0.03 | 6.7 ± 0.5 | 97 ± 1 | 15 ± 1 |
| A347G | 5.8 ± 0.03 | 5.8 ± 0.3 | 1.0 ± 0.1 | 7.9 ± 0.4 | 133 ± 5 | 17 ± 1 |
| L353A | 6.7 ± 0.03 | 1.9 ± 0.1 | 0.3 ± 0.01 | 6.9 ± 0.2 | 57 ± 0.7 | 8.4 ± 0.2 |
| R354A | 7.9 ± 0.1 | 2.7 ± 0.04 | 0.3 ± 0.01 | 8.0 ± 0.3 | 75 ± 2 | 9.3 ± 0.4 |
| V362A | 5.6 ± 0.1 | 2.4 ± 0.02 | 0.4 ± 0.01 | 5.7 ± 0.1 | 47 ± 0.7 | 8.2 ± 0.2 |
| N363A | 6.8 ± 0.1 | 4.0 ± 0.1 | 0.6 ± 0.02 | 8.1 ± 0.3 | 89 ± 2 | 11 ± 0.6 |
| H372A | 5.3 ± 0.1 | 35 ± 0.4 | 6.5 ± 0.1 | 7.5 ± 0.5 | 78 ± 6 | 10 ± 1 |
| A375G | 5.1 ± 0.1 | 6.1 ± 0.1 | 1.2 ± 0.03 | 6.2 ± 0.2 | 123 ± 5 | 20 ± 1 |
| A376G | 7.2 ± 0.03 | 10 ± 0.4 | 1.4 ± 0.1 | 8.2 ± 0.3 | 121 ± 4 | 15 ± 0.7 |
| I377A | 6.9 ± 0.04 | 2.1 ± 0.01 | 0.3 ± 0.002 | 7.3 ± 0.1 | 68 ± 0.5 | 9.3 ± 0.2 |
| A378G | 6.5 ± 0.1 | 1.5 ± 0.01 | 0.2 ± 0.002 | 7.2 ± 0.1 | 45 ± 0.8 | 6.2 ± 0.1 |
| K380A | 5.4 ± 0.03 | 5.1 ± 0.1 | 0.9 ± 0.02 | 6.3 ± 0.4 | 112 ± 4 | 18 ± 1 |
| A382G | 6.9 ± 0.04 | 1.0 ± 0.01 | 0.1 ± 0.001 | 6.9 ± 0.1 | 35 ± 0.1 | 5.0 ± 0.1 |
| V386A | 6.6 ± 0.04 | 1.9 ± 0.1 | 0.3 ± 0.01 | 6.5 ± 0.1 | 53 ± 0.3 | 8.2 ± 0.2 |
| I388V | 7.2 ± 0.1 | 3.3 ± 0.04 | 0.5 ± 0.01 | 7.5 ± 0.2 | 85 ± 0.8 | 11 ± 0.3 |
| R399A | 8.0 ± 0.1 | 7.3 ± 0.1 | 0.9 ± 0.01 | 9.2 ± 1 | 193 ± 14 | 21 ± 3 |
| F400A | 7.5 ± 0.5 | 110 ± 4 | 15 ± 1 |  |  |  |
| E401A | 7.3 ± 0.1 | 5.3 ± 0.03 | 0.7 ± 0.01 | 7.8 ± 0.5 | 163 ± 6 | 21 ± 2 |

**Supplementary Table 5.** *k*_*on*_, *k*_*off*_ and *K*_*d*_ for the interaction between PSG and CRIPT 15 AA or CRIPT _P-3_ 15 AA.

|  | GKKVLDTKNYKQTSV |  |  | GKKVLDTKNYKNvaTSV |  |  |
| --- | --- | --- | --- | --- | --- | --- |
| | $k_{on}$ (s<sup>-1</sup> μM<sup>-1</sup>) | $k_{off}$ (s<sup>-1</sup>) | $K_d$ (μM) | $k_{on}$ (s<sup>-1</sup> μM<sup>-1</sup>) | $k_{off}$ (s<sup>-1</sup>) | $K_d$ (μM) |
| PSG | 8.2 ± 0.1 | 3.5 ± 0.02 | 0.43 ± 0.01 | 7.0 ± 0.2 | 12 ± 0.1 | 1.7 ± 0.1 |
| G322A | 8.2 ± 0.1 | 0.8 ± 0.01 | 0.10 ± 0.002 | 7.0 ± 0.1 | 2.4 ± 0.03 | 0.34 ± 0.01 |
| F325A | 6.2 ± 0.2 | 8.8 ± 0.1 | 1.41 ± 0.05 | 6.1 ± 0.3 | 22 ± 0.2 | 3.6 ± 0.2 |
| I327V | 6.3 ± 0.2 | 19 ± 0.6 | 3.0 ± 0.1 | 5.6 ± 0.3 | 52 ± 0.1 | 9.2 ± 0.5 |
| G330A | 7.7 ± 0.3 | 99 ± 2.6 | 13 ± 0.7 | 15 ± 2 | 208 ± 13 | 14 ± 2 |
| E334A | 9.4 ± 0.1 | 8.4 ± 0.2 | 0.90 ± 0.02 | 8.2 ± 0.3 | 28 ± 0.5 | 3.4 ± 0.1 |
| G335A | 8.3 ± 0.1 | 1.8 ± 0.03 | 0.21 ± 0.005 | 7.2 ± 0.1 | 5.3 ± 0.1 | 0.73 ± 0.02 |
| I338A | 7.1 ± 0.1 | 2.7 ± 0.01 | 0.38 ± 0.004 | 6.2 ± 0.1 | 8.2 ± 0.04 | 1.3 ± 0.02 |
| A347G | 4.8 ± 0.1 | 8.9 ± 0.4 | 1.9 ± 0.1 | 2.9 ± 0.2 | 35 ± 0.7 | 12 ± 1 |
| L353A | 8.7 ± 0.1 | 2.8 ± 0.01 | 0.32 ± 0.005 | 7.4 ± 0.2 | 8.9 ± 0.1 | 1.2 ± 0.03 |
| R354A | 8.1 ± 0.2 | 3.9 ± 0.03 | 0.48 ± 0.01 | 7.8 ± 0.2 | 13 ± 0.2 | 1.6 ± 0.04 |
| V362A | 4.7 ± 0.3 | 3.6 ± 0.2 | 0.78 ± 0.1 | 6.5 ± 0.1 | 12 ± 0.1 | 1.9 ± 0.03 |
| A376G | 8.1 ± 0.1 | 12 ± 0.1 | 1.4 ± 0.02 | 5.4 ± 0.2 | 28 ± 0.3 | 5.1 ± 0.2 |
| R399A | 8.1 ± 0.2 | 5.0 ± 0.1 | 0.62 ± 0.02 | 8.0 ± 0.1 | 15 ± 0.1 | 1.8 ± 0.03 |
| F400A | 6.3 ± 0.1 | 6.4 ± 0.1 | 1.0 ± 0.02 | 6.9 ± 0.3 | 41 ± 1 | 6.0 ± 0.3 |
| E401A | 6.8 ± 0.1 | 3.4 ± 0.04 | 0.50 ± 0.01 | 6.4 ± 0.1 | 12 ± 0.1 | 1.9 ± 0.04 |

**Supplementary Table 6.** *k*_*on*_, *k*_*off*_ and *K*_*d*_ for the interaction between PSG and CRIPT 6 AA or CRIPT _P-3_ 6 AA.

|  | YKQTSV |  |  | YKNvaTSV |  |  |
| --- | --- | --- | --- | --- | --- | --- |
| | $k_{on}$ (s<sup>-1</sup> μM<sup>-1</sup>) | $k_{off}$ (s<sup>-1</sup>) | $K_d$ (μM) | $k_{on}$ (s<sup>-1</sup> μM<sup>-1</sup>) | $k_{off}$ (s<sup>-1</sup>) | $K_d$ (μM) |
| PSG | 6.0 ± 0.1 | 1.3 ± 0.1 | 0.21 ± 0.02 | 5.8 ± 0.1 | 2.1 ± 0.2 | 0.4 ± 0.03 |
| G322A | 6.8 ± 0.1 | 0.26 ± 0.002 | 0.04 ± 0.001 | 6.0 ± 0.1 | 1.0 ± 0.1 | 0.2 ± 0.01 |
| F325A | 5.5 ± 0.2 | 4.7 ± 0.5 | 0.87 ± 0.1 | 5.2 ± 0.3 | 8.3 ± 0.3 | 1.6 ± 0.1 |
| I327V | 5.7 ± 0.1 | 8.9 ± 1.0 | 1.6 ± 0.2 | 4.4 ± 0.2 | 15 ± 0.1 | 3.4 ± 0.1 |
| G330A | 6.2 ± 0.2 | 35 ± 1.8 | 5.8 ± 0.3 | 6.6 ± 0.2 | 85 ± 2.6 | 13 ± 0.6 |
| E334A | 7.2 ± 0.1 | 2.4 ± 0.1 | 0.33 ± 0.01 | 6.4 ± 0.2 | 6.8 ± 0.1 | 1.1 ± 0.04 |
| G335A | 7.0 ± 0.1 | 0.74 ± 0.03 | 0.11 ± 0.01 | 6.9 ± 0.2 | 1.8 ± 0.03 | 0.3 ± 0.01 |
| I338A | 5.4 ± 0.1 | 1.1 ± 0.03 | 0.20 ± 0.01 | 4.7 ± 0.1 | 2.7 ± 0.2 | 0.6 ± 0.04 |
| A347G | 4.4 ± 0.1 | 5.0 ± 0.3 | 1.1 ± 0.1 | 3.8 ± 0.1 | 14 ± 0.6 | 3.6 ± 0.2 |
| L353A | 6.3 ± 0.1 | 1.1 ± 0.1 | 0.18 ± 0.01 | 5.6 ± 0.1 | 3.3 ± 0.2 | 0.6 ± 0.03 |
| R354A | 7.0 ± 0.1 | 1.7 ± 0.1 | 0.25 ± 0.02 | 5.9 ± 0.1 | 4.7 ± 0.1 | 0.8 ± 0.03 |
| V362A | 7.1 ± 0.4 | 1.3 ± 0.01 | 0.19 ± 0.01 | 5.4 ± 0.3 | 6.8 ± 0.9 | 1.3 ± 0.2 |
| A376G | 6.6 ± 0.1 | 5.4 ± 0.2 | 0.82 ± 0.03 | 5.6 ± 0.2 | 14 ± 1.2 | 2.4 ± 0.2 |
| R399A | 7.2 ± 0.2 | 2.8 ± 0.3 | 0.38 ± 0.04 | 6.3 ± 0.2 | 6.7 ± 0.4 | 1.1 ± 0.1 |
| F400A | 6.4 ± 0.3 | 13 ± 0.4 | 2.0 ± 0.1 | 4.0 ± 0.1 | 29 ± 0.4 | 7.2 ± 0.2 |
| E401A | 5.6 ± 0.1 | 5.1 ± 0.5 | 0.90 ± 0.1 | 5.1 ± 0.3 | 7.3 ± 0.4 | 1.4 ± 0.1 |

**Supplementary Table 7.**
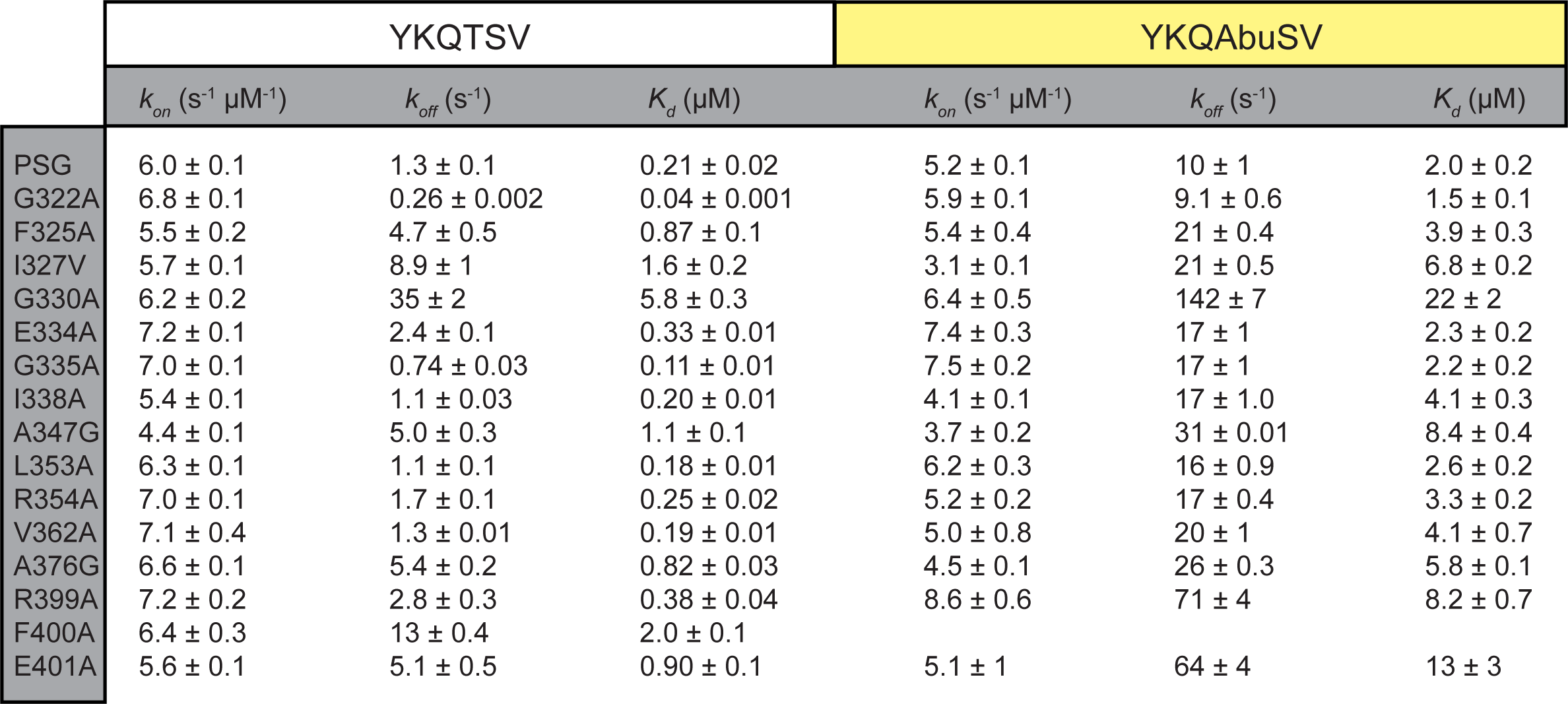
*k*_*on*_, *k*_*off*_ and *K*_*d*_ for the interaction between PSG and CRIPT 6 AA or CRIPT _P-2_ 6 AA

